# The representation and valuation of subgoals in the human brain during model-based hierarchical behavior

**DOI:** 10.1101/2025.03.24.645084

**Authors:** Cooper D. Grossman, Vincent Man, John P. O’Doherty

## Abstract

The human capacity to plan and perform long, complex sequences of behavior to achieve distant goals depends in part on a hierarchical organization that divides behavior into structured segments. Such a mechanism requires the internal designation of certain states as subgoals to mark the successful implementation of a behavioral segment. How the brain represents subgoals over time and computes decision values as a function of subgoals is unknown. While most characterizations of hierarchical behavior lack knowledge of the environment, human decision-making also relies on planning with an internal model of the world. Consequently, it remains to be determined how the brain computes values of subgoals using model-based planning in order to drive hierarchical, model-based decisions. Using a sequential-subgoal decision-making task designed to evoke hierarchical, model-based behavior in combination with fMRI, we decoded a representation of the current subgoal in insula and ventromedial prefrontal cortex during decision-making that persisted over time–a critical, latent representation for computing values and orienting behavior in the correct sequence. Using a model-based, hierarchical reinforcement learning model, we also found key decision signals based on values from the model in several regions of frontal cortex. These findings thereby shed light on the neural correlates of subgoal representation and illustrate how value signals can be computed on the basis of these subgoals and knowledge of the environment structure.

---

Human decision-making often requires complicated and temporally-extended sequences of behavior to achieve goals. To make these goals more tractable, such complex tasks can be broken down into different components, each with their own “subgoals” (Sutton et al., 1999, sometimes called “pseudo-rewards”). Subgoals are internally designated states of the world that a person can choose to pursue as an interim means of progressing toward an overall goal. The brain can make decisions over these subgoals and then evaluate choices relative to a chosen subgoal. The use of subgoals instantiates a hierarchy in cognition that allows for more efficient decision making through segmentation, abstraction, and generalization (Lashley, 1951; Miller et al., 1960; Norman and Shallice, 1986; Hengst, 2010). Subgoals allow for simpler planning by breaking up the sequence of behavior. Reinforced by subgoals, sequences of actions can be compressed and treated as abstract “options” during decision making. These options and their subgoals can then be used across different goals, facilitating generalization of behavior. Since subgoals are outcomes designated as reinforcing by the brain, they are distinct from the well-studied primary reinforcers such as food and water, or strong secondary reinforcers such as money (hereafter, “external reinforcers”). While previous neural and behavior evidence substantiates the theoretical notion of subgoals, how the brain represents subgoals over behaviorally relevant timescales and computes values as a function of them is still unknown. Given a goal, the brain must maintain a representation of the appropriate subgoal in order to orient behavior towards that subgoal. The ventromedial prefrontal cortex (vmPFC) has consistently been implicated in representing goals and goal values (O’Doherty, 2011). This region has been implicated in signaling value that generalizes across goals (Plassmann et al., 2007; Hare et al., 2008; McNamee et al., 2013) but can also represent and discriminate between goal identities (Levy and Glimcher, 2011; McNamee et al., 2013). However, all of these signals refer to external reinforcers. As a result, a fundamental question remains as to how the brain, and the vmPFC in particular, represent subgoals during hierarchical behavior (Bein and Niv, 2023). It is also unclear how these representations persist over time to drive ongoing behavior, given that a sustained representation of a given subgoal would likely be necessary to ensure that decisions remain focused on attaining that subgoal.

Broadly, previous research supports the notion that the brain uses hierarchical representations (Badre and Nee, 2018). Research focusing specifically on hierarchical processes that include action sequences oriented towards subgoals has indicated a special role for medial frontal cortex (mFC), particularly in the anterior cingulate cortex and more dorsal. This region has been found to represent learning signals related to subgoals (Ribas-Fernandes et al., 2011, 2019) and respond to states that are theoretically likely to be designated as subgoals (Balaguer et al., 2016). The region is involved in other aspects of hierarchical cognition, like abstract plan complexity (Balaguer et al., 2016) and in distinguishing different abstract task states (Holroyd et al., 2018; Colin et al., 2023).

One gap in this literature is how the brain computes values in relation to subgoals—a crucial component of hierarchical control. A wealth of previous studies in humans has implicated both the dorsal medial frontal cortex (dmFC; Wunderlich et al., 2009; Lee et al., 2014; Kim et al., 2019; Aquino et al., 2023) and medial prefrontal cortex (mPFC; for review see Clithero and Rangel, 2013) in computing decision values as a function of external reinforcers, but not subgoals. Holroyd and colleagues have theorized that a specific region of mPFC, rostral anterior cingulate cortex (rACC), is involved in hierarchical task representation and decision-making, but this theory has not been explicitly tested (Holroyd and Yeung, 2012; Holroyd and Verguts, 2021; Alejandro and Holroyd, 2024). Consequently, it remains unclear whether or how decision values related to the pursuit of subgoals are represented in these regions or elsewhere in the brain.

Hierarchical behavior has often been modeled using model-free reinforcement learning (RL), in which agents learn only through direct experience with goals and subgoals (Botvinick, 2012; Rasmussen et al., 2017; Eckstein and Collins, 2020; Xia and Collins, 2021; Li et al., 2022). Without understanding the environment’s structure, this approach cannot account for latent learning: learning about outcomes that are not relevant to the current policy but can be used for future behavior. This model-free framework also faces the “option discovery problem” of identifying states of the world as useful subgoals (Sutton et al., 1999; Precup, 2000). In contrast, model-based RL captures human behavior more effectively in this regard, as it involves building an internal model of the environment to prospectively plan and compute action values based on state transitions (for review see Collins and Cockburn, 2020; Drummond and Niv, 2020). This structure enables latent learning and could facilitate option discovery (Botvinick and Weinstein, 2014). On the other hand, model-based RL becomes computationally intractable when considering all possible state transitions and relies on a costly re-computing of action values at each step. Additionally, it remains unclear which perceptual and semantic details should be included in state representations to ensure efficiency and relevance of the internal model. Integrating hierarchical processes into model-based frameworks may resolve these inefficiencies by compressing state-action space and reducing the number of value computations performed (Botvinick and Weinstein, 2014; Alejandro and Holroyd, 2024). Off-policy subgoals could also guide latent learning, resulting in an internal model that includes the most behaviorally relevant information. The cooperative integration of model-based and hierarchical processes has received attention in machine learning and some in neuroscience, but many questions remain about their intersection (Botvinick and Weinstein, 2014; Balaguer et al., 2016; Chalmers et al., 2016; Tomov et al., 2020; Pateria et al., 2021). One theory that has arisen in the context of explaining human behavior is “saltatory” model-based hierarchical RL (MB-HRL; Botvinick and Weinstein, 2014). MB-HRL executes model-based planning using options that jump over parts of state space in its internal model. How the model would construct options, which assume a certain path through a probabilistic state space, is still unknown. And while a Bayesian version of this model has shown utility in capturing hierarchy discovery (Tomov et al., 2020), the MB-HRL theory has not been directly tested in decision-making behavior. Thus, the potential behavioral and neural mechanisms of MB-HRL are entirely unknown.

Here, we designed a novel sequential subgoal decision-making task to better understand how the brain represents subgoals and values computed in relation to those subgoals. The task was simultaneously designed to better understand questions at the intersection of hierarchical and model-based cognition in human brain and behavior, like how options are learned in stochastic environments. Participants performing the task in an fMRI scanner exhibited behavior that was consistent with both model-based and hierarchical processes, making decisions that reflected subgoal-driven latent learning to efficiently collect subgoals in a stochastic environment. Response times indicated that subjects planned sequences of actions over probabilistic state transitions. These planned actions reflected an assumption of a common state transition, exploiting the statistical regularities of the environment. Substantiating the notion that subjects planned actions over probabilistic transitions, actions could also be decoded from motor cortex and amygdala prior to the transition. We adapted an MB-HRL model that was sufficient to recapitulate several aspects of behavior. Crucial to orienting hierarchical behavior in the correct sequence, a latent representation of the current subgoal was decoded in insula and vmPFC at the initiation of choices that persisted over the course of the trial. Estimated from this model, decision variables that integrate values—computed as a function of subgoals and using knowledge of the environment—were found to be represented in multiple regions of mFC. Thus, we substantiate a framework for hierarchical, model-based cognition, providing neural and behavioral evidence for a strategy that computes values as a function of subgoals and knowledge of environment structure in order to make decisions.

## Results

### The sequential subgoal decision-making task tests for model-based, hierarchical behavior

To examine how the brain represents and makes decisions about subgoals, we developed a sequential subgoal task for human participants (Figure 1A,B). In this task, subjects make decisions to complete a sequence of 4 subgoals, in the appropriate order, to obtain a monetary goal. The task is presented as a “Space Taxi” game in which the participant navigates their taxi to different planets in order to sequentially acquire the subgoals resulting in the collection of a monetary fare. The subgoals are collecting a permit, picking up two aliens (one at a time), and then dropping those aliens off at their home planet. The planets are the terminal states of a two-step task structure that requires two sequential choices (left or right) in order to reach one of those states and complete the trial (Gläscher et al., 2010; Lee et al., 2014; Weissengruber et al., 2019; Kim et al., 2019). The subject starts at Earth (response state 1) on every trial and makes a choice between a left and a right portal. Each choice leads to one of two space stations, with high (*P* = 0.7) or low (*P* = 0.3) probability (response state 2). The stochasticity in this environment allows for the assessment of model-based decision-making that considers this structure. A second left or right portal choice at the space station leads deterministically to one of four planets (outcome state) containing one of the subgoals or nothing. The subject may encounter any of the subgoals, but can only complete it in the correct order. For example, if the subject needs a permit, they may find an alien but cannot pick it up. Whether or not the subgoal was completed is indicated to the subjects following the outcome state. This design permits examination of another key feature of model-based cognition: latent learning about the structure of an environment that can be exploited later on. The relationship between Earth and the space stations stays the same, but with each new fare the planets are pseudo-randomized (ensuring all subgoals can be found by common transitions). This design forces subjects to learn the new state-action space with each new fare, facilitating the examination of latent learning. In order to confirm that subjects maintained a correct understanding and memory of subgoal sequence, randomly (*P* = 1*/*3) on some trials subjects were probed to report which subgoal they intended to pursue (see Methods). The monetary value of the fare also varied across fares ($0.10, $0.30, or $1.00) and was indicated to the subjects at the beginning of each new fare.

**Figure 1.**
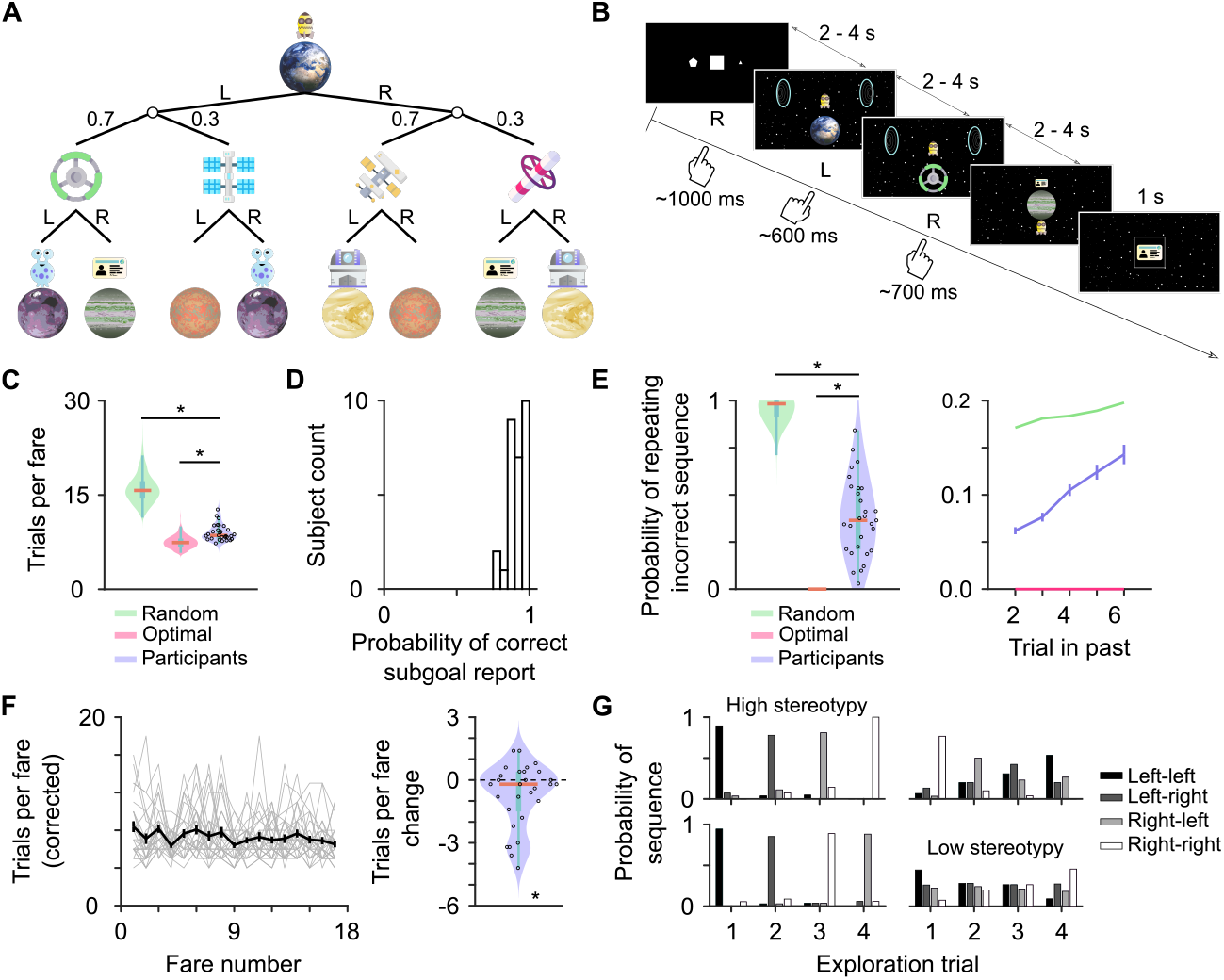
Subjects make hierarchical decisions to collect subgoals and complete fares. (A) Schematic of task state structure for an example fare. (B) Example trial from the perspective of the subject. (C) Mean trials per fare for an agent that selects actions randomly, an optimal model-based agent with perfect learning and memory, and the participants. Top and bottom edges of blue boxes represent the 75th and 25th percentile, respectively. Orange line is the median of the data. Subject performance was significantly better than the random agent (*W* = 5.29 *×* 10^5^, *p <* 10^*−*19^, Wilcoxon rank sum test) but worse than the optimal agent (*W* = 5.01 *×* 10^5^, *p <* 10^*−*224^, Wilcoxon rank sum test). (D) Fraction of trials on which the participants correctly report which subgoal is the current one. (E) Left: the probability of making a sequence of choices that is incorrect based on all previous choices. Limited to common transition trials. Subject performance was significantly better than the random agent (*W* = 5.3 *×* 10^5^, *p <* 10^*−*19^, Wilcoxon rank sum test) but worse than the optimal agent (*W* = 5.0 *×* 10^5^, *p <* 10^*−*224^, Wilcoxon rank sum test) Right: the probability of repeating incorrect choices increased as a function of how far in the past those choices were last made (effect of trial in past, coefficient = 0.14, *t*_14628_ = 7.16, *p <* 10^*−*12^, linear mixed-effects model). (F) Left: Number of trials per fare by fare number within session. To correct for stochasticity in the task, rare transition trials without subgoal collection were subtracted from values. Plot limited to the minimum number of fares completed by any participant. Errorbars represent SEM. Right: differences in performance between first and last 5 fares of each participant’s session (mean trials per fare in last 5 fares - mean trials per fare in the first 5 fares) demonstrating an average improvement in performance (effect of fare number: coefficient = *−*0.048, *t*_760_ = *−*4.00, *p <* 10^*−*4^, linear mixed-effects model). (G) Exploration patterns for 4 example subjects with varying patterns and amounts of stereotypy. Exploration trials occur when the subject does not have previous experience with the current subgoal. Limited to common transition branches of the state-action space, there are 4 possible exploration trials and sequences. Top left and bottom left subjects represent the most common patterns of exploration with high stereotypy.

### Sequential choices reflect hierarchical, model-based behavior

Participants (n = 29) performed the Space Taxi game while undergoing functional magnetic resonance imaging (fMRI). Subjects completed 222 (3 blocks, n = 3) or 262 trials (4 blocks, n = 26) during the session. The included subjects completed fares efficiently (Figure 1C; 9.01 ± 1.29 trials per fare). The average number of trials that it took to complete a fare was closer to the performance of an optimal agent than a random one (Figure 11C, Figure S1A,B). This performance was not affected by the fare value (Figure S1A) and nor were first state response times (*p* = 0.93, linear mixed-effects model). Performance was partially driven by the appropriate execution of the subgoal sequence; subjects consistently tracked the correct order of subgoal completion, correctly reporting the current subgoal with high reliability (probability correct, 0.91 ± 0.059, Figure 1D). Errors in this measure could mainly be attributed to the difficulty of the reporting mechanism (Figure S1D; see Methods). We also assessed choices on the first trial of a new subgoal (excluding second alien), after collecting the previous subgoal. If subjects did not understand the sequential structure then they would make the same choices as the previous trial at chance level. We found that subjects very rarely (*P* = 0.053 ± 0.057) repeated their choices after collecting a subgoal, significantly below chance (*P* = 0.25 with 4 possible choice sequences; one-sample *t*-test, *t*_28_ = −18.51, *p* = 0). These results demonstrate that subjects understood the sequential task structure and made decisions hierarchically, contingent upon correct knowledge of the current subgoal.

Suboptimality in behavior was found to be a consequence of subjects making incorrect sequences of choices— choices that had been previously made that do not lead to the current subgoal or failing to make choices that were previously made and do lead to the current subgoal (Figure 1E). The rate of incorrect choices increased as those choices last occurred further back in time (Figure 1E). This pattern of lapses suggests that the state-action space was forgotten over time within a fare. We also found that performance improved over the course of the session for some subjects. The number of trials it took to complete a fare tended to be worse at the beginning of the session than at the end (Figure 1F) which was also the case for the probability of repeating an incorrect sequence (Figure S1E). Response times in the first state also sped up over the course of the session (effect of trial number in session, *t*_7247_ = −5.99, *p <* 10^*−*8^, linear mixed-effects model). Unexpectedly, we also found that many subjects exhibited stereotyped patterns of exploration, exploring the different parts of the state-action space in the same order on trials when there was no previous experience with the location of the current subgoal (Figure 1G, Figure S1F). Interestingly, greater exploration stereotypy predicted better performance in the task (Figure S1G).

To demonstrate that behavior in this task is model-based, we examined indicators that decisions leveraged knowledge of the sequential task structure and the stochastic transition structure of the environment. A fundamental feature of model-based behavior is the learning of the state-action space, which may not be immediately apparent in behavior but can be exploited later on. If subjects are engaged in latent learning about the state-action space, then choices should be correct on the first trial of a new subgoal if the subject has encountered that subgoal previously (while it was *not* the current subgoal). We found that previous experience conferred a significant advantage to the probability of making the correct sequence of actions, compared to instances in which there was no previous experience (Figure 2A). Even without previous experience latent learning should provide some advantage, since model-based processes will also learn where the subgoal *cannot* be found. Indeed, when compared to a model-based agent that forgets its knowledge of the environment with each new subgoal (no off-subgoal learning), the subjects show greater probability of performing the correct sequence of actions on the first trial of a new subgoal without previous experience (Figure 2A). The use of this knowledge is also implicit in the observed exploration patterns of some subjects; because exploration trials are defined by not having previous experience with the subgoal, the patterns in choice probabilities demonstrate that subjects are not repeating choice sequences that will not lead to the current subgoal (Figure 1G).

**Figure 2.**
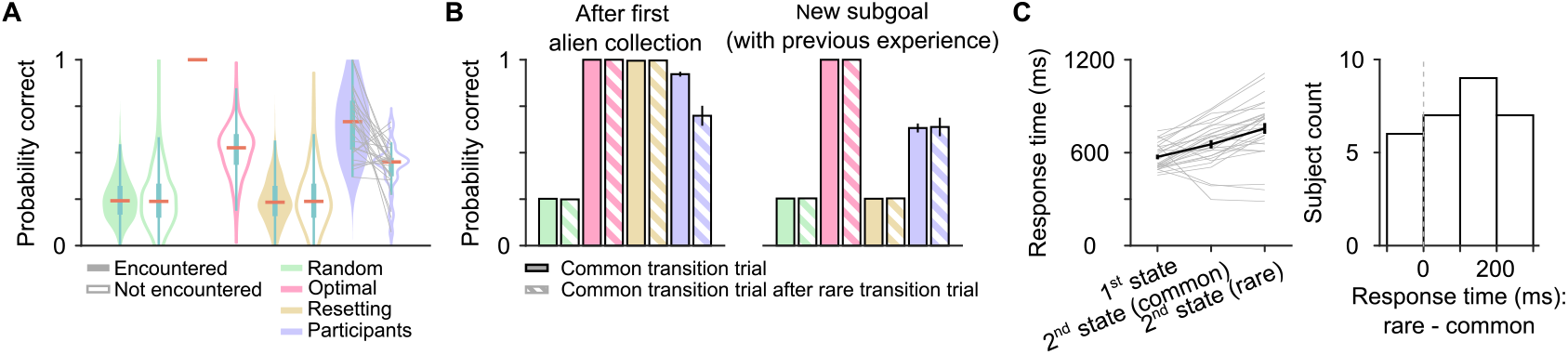
Subjects make choices based on the task structure. (A) Probability of making the correct sequence of choices on the first trial of a new subgoal, split by whether or not that subgoal was encountered previously or not. Limited to common transition trials. Actual subjects (magenta), compared to random (cyan), optimal (blue), and resetting (purple) agents. Top and bottom edges of blue boxes represent the 75th and 25th percentile, respectively. Orange line is the median of the data. Gray lines indicate probabilities for individual subjects across conditions. Subject choices were more often correct when there was a previous encounter with the subgoal (*t*_28_ = 5.21, *p <* 10^*−*4^, paired t-test). (B) Left: Probability of making the correct sequence of actions on the last trial of different sequences of common and rare transition trials when pursuing the second alien. Right: Probability of making the correct sequence of actions on the first common transition trial of a new subgoal when there was previous experience with that subgoal. Some of these precise sequences are rare, so individual data are pooled before averaging. Colors indicate same groups as in A. Errorbars represent Bernoulli SEM. (C) Left: response times by task state and transition type. Errorbars represent SEM. Right: differences in second state response times between rare and common transition reveal an effect of transition type (effect of rare transition *t*_7244_ = 6.18, *p <* 10^*−*9^, linear mixed-effects model).

Model-based decisions should also take into account knowledge of probabilistic state transitions. To examine this possibility we assessed the affect of rare transitions when subjects could have had knowledge of the correct sequence of actions. In one case, we looked at instances in which subjects collected their first alien on a common transition, experienced a rare transition trial without alien collection, and experienced a common transition on the third trial. The correct choice probability was compared to trials after collection of the first alien without the intervening rare transition trial. We also examined the second trial of a new subgoal when there was previous experience (potential latent learning) and the first trial was a rare transition trial. Similar to the behavioral signatures described in previous studies (Gläscher et al., 2010; Daw et al., 2011), purely model-based behavior should understand the environment structure and be unaffected by the outcome of the intervening rare transition trial. In both instances, we found that subjects continued with the correct sequence of actions on the next trial with a common transition with high probability (Figure 2B). Between aliens however, there was a decrease in accuracy that could be attributed to forgetting but may also be the result of additional contributions from a model-free learner (see Discussion). We also found that second choices on the intervening rare transition trials were different from the correct choices on the subsequent common transition trial, demonstrating that subjects were not ignoring the identity of the second response state (Figure S2A). Similarly, choices on the first trial of a new subgoal reflected knowledge of the probabilistic state transitions (Figure S2B). As described above, choices on these trials rarely matched choices on the previous collection trial when the transition type (common or rare) was the same. However, when transition types were different, the probability of repeating the same sequence of choices was near chance (*P* = 0.21 ± 0.14, Figure S2C). These choice patterns demonstrate that the participants understand the probabilistic state structure.

Subjects also exhibited behavior consistent with the use of options (hierarchy over actions). Response times were slower on response state 2 following a rare transition than after a common one (Figure 2C). This difference suggests that subjects had an expectation about which state is coming next based on statistical regularities in the state-action space. It may also indicate that subjects had already planned their second action based on this expectation. This finding is consistent with observations that humans learn statistical spatiotemporal regularities (Hunt and Aslin, 2001; Fiser and Aslin, 2002), learn higher-order environmental structure (Schapiro et al., 2013; Kahn et al., 2018), and simplify that structure (Lynn et al., 2020) to facilitate faster responses. A bigger difference between response times after rare or common transitions was also found to predict better performance in the task (Figure S2C). This finding is consistent with the idea that better learners of task structure are more likely to plan sequences that exploit that structure.

### MB-HRL provides a computational framework for behavior

To understand the potential cognitive mechanisms that produce this model-based, hierarchical behavior, we designed a generative model of behavior based on an existing theory of MB-HRL (“saltatory MB-HRL”; Botvinick and Weinstein, 2014). In the same manner as previous model-based approaches, this agent learns an internal model of state-action space by learning the transition probabilities between states of the task, contingent upon taken actions (Gläscher et al., 2010; Daw et al., 2011). Importantly, the terminal outcome state representations contain information about subgoal identity. While it is not known what information is generally included in the representation of a state, this information should be behaviorally relevant. Previous research has also shown behavior driven by state representations that contain information about goal identity (Kim et al., 2019).

At a more abstract level of state representation, we assume the agent has already learned the *task* state space model—essentially, a separate internal model that defines the transition probabilities between the different subgoals in the sequence. This assumption is justified by the explicit instructions given to the subjects about this aspect of task structure, reliability of the subgoal report, and the other behavior metrics assessed above that demonstrate participant comprehension of task structure. Using this task state model, the agent tracks the current subgoal and updates this representation in the correct order after each subgoal is collected (i.e., deterministically transitioning between current subgoal states). The agent uses the representation of current subgoal as context for valuation. The current subgoal determines values of the outcome states and subsequently the agent can use the learned state-action space to compute decision values through prospective planning.

At a more behaviorally fine-grained level of hierarchy, the agent abstracts over sequences of two actions and decides between these options during the first response state. These options rely on a type of internal option model, similar to the state-action model, that abstracts over the observable states of the task (Botvinick and Weinstein, 2014). Unique to our design, the option model assumes a common transition between the first and second response states. Thus, in the first state, the model decides between four possible options (left-left, left-right, right-left, and right-right). This design implements one possible solution to the problem of how options and option models may be formed in probabilistic environments: options are formed across transitions in the environment (represented by the internal model of state-action space) that have higher statistical regularity. The abstraction of options and option model simplifies planning and avoids the requirement of re-computing values after every state transition. The options model also contains a termination function that determines when the option should end (Botvinick and Weinstein, 2014). In this task, the option ends after the outcome state or when it is interrupted by a rare transition. When the option is terminated after a rare transition, the values of the two available actions are computed to make a decision.

On top of this central framework, we added several parameters to capture other features of behavior. Exploration biases (similar to novelty bonuses; Kakade and Dayan, 2002; Wittmann et al., 2008; Krebs et al., 2009; Cockburn et al., 2022) were added into the decision function to capture stereotyped exploration patterns (Figure 1G). Each bias multiplied the computed value for an option until that option was explored. To capture forgetting of the state-action space (Figure 1E), learned transition probabilities were forgotten over time. Since this forgetting improved over the course of the session for some subjects (Figure 1F), forgetting was controlled by an initial rate that could change linearly as a function of trial number in the session.

### MB-HRL recapitulates multiple facets of behavior

Our implementation of the full MB-HRL model described above performed better than versions lacking any of the individual components as well as a “resetting” version that forgets the state-action space with each new subgoal (Figure S3A). The full model was also sufficient to reproduce overall performance (Figure 3A) and several key elements of behavior. Behavior simulations using fit parameters accurately recapitulated patterns of latent learning (Figure 3B) and the forgetting of it (Figure S3b). Simulations also demonstrated the same responses to the transition structure on trials in between first and second aliens and on the first trial of a new subgoal when latent learning could have occurred (Figure 3C). The decrease in accuracy between aliens when there was a rare trial in between, and forgetting patterns generally, were also recapitulated by model simulations (Figure 3C,D). The parameterized change in forgetting reproduced the improvement in behavior over the course of the session (Figure 3E and Figure S3C). Individual patterns of exploration matching those of the subjects were also observed in simulated behavior (Figure 3F). Simulations using models that lacked one feature of the full model were unable to capture all features of behavior (Figure 3A-F). The subject-level parameters of the behavior model also captured individual variability, predicting observed performance, improvement, and exploration stereotypy (Figure S3D).

**Figure 3.**
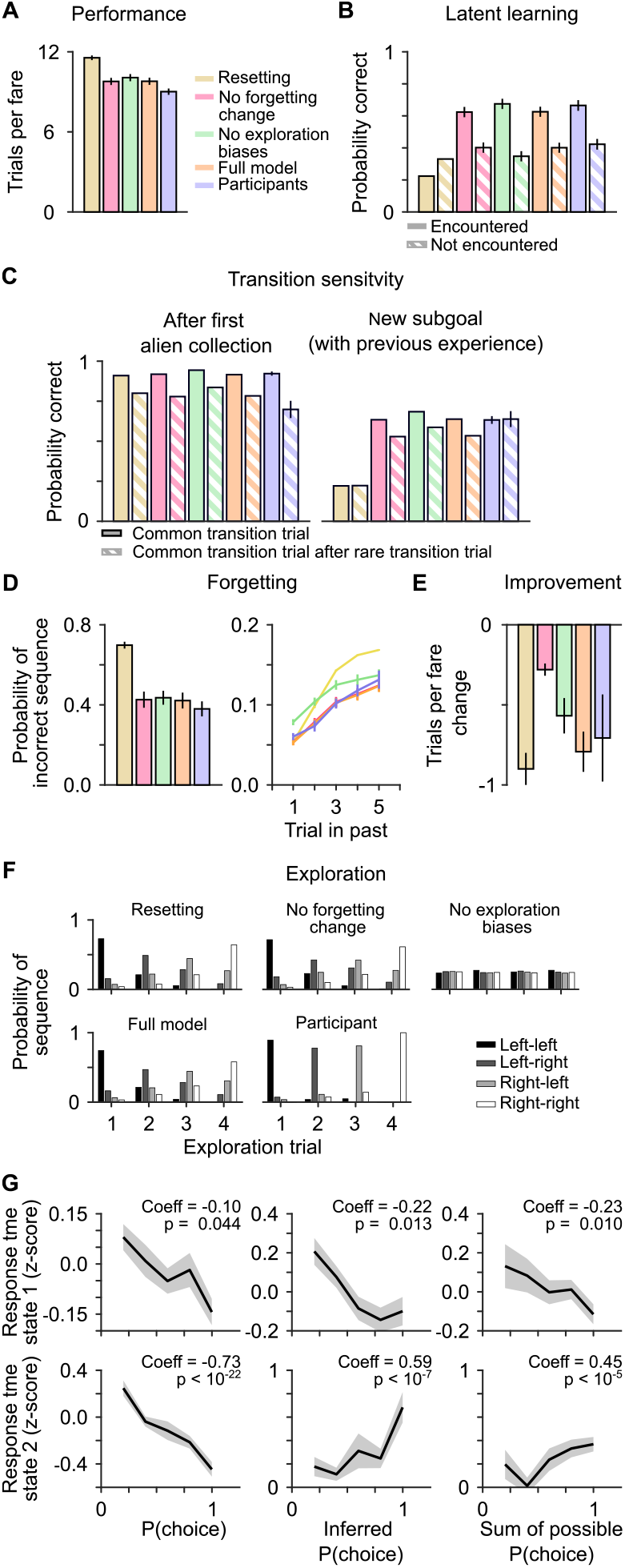
MB-HRL recapitulates behavior. Actual behavior from participants compared to behavior simulated from the fitted, full MB-HRL model and models lacking one of the features of the full model. Note that some of the colors indicate different models from the previous figures. (A) Overall performance as in Figure 1c. (B) Expression of latent learning as in Figure 2a. (C) Signatures of model-based behavior in response to the probabilistic task structure, as in Figure 2b. (D) Forgetting behavior, as in Figure 1c. (E) Behavior improvement over the session, as in Figure 1f. (F) Exploration biases from one example subject, as in top left panel of Figure 1g. (G) Response times (z-scored within subject) as a function of MB-HRL choice probability metric, task state, and transition type. Coefficients and *p* values from linear mixed-effects models.

The difference in response times during the second response state as a function of transition type that we observed may have come as result of an interruption in action preparation involved in sequential behavior or simply as a consequence of the surprising event itself. This distinction is important for distinguishing a model that decides over options from one that decides over single actions. To further substantiate the model framework that uses options, we examined the relationship between the probability of the chosen option computed by the model and response times. This decision variable represents the exploration-weighted value of an option relative to the others. This weighted value integrates value computed as a function of the learned state-action space as well as the exploration biases. This choice probability is the model output that drives decisions and may also correspond to decision confidence. We first compared the chosen option probability to response times on common transition trials, when the chosen option was known (on rare transition trials an option planned assuming a common transition cannot be observed). We found that higher option probabilities predicted faster responses on both choices in the sequence, revealing a linear relationship (Figure 3G). On rare transition trials, we computed two choice probability terms as a proxy for chosen option probability. First, we summed the probabilities of the two possible options that could have been taken, given the observed first choice. Second, we inferred the chosen option by finding the maximum option probability of the two possible. These terms predicted future behavior on the next common transition trial (Figure S3E). For both terms, increasing values predicted faster response times in the first response state (Figure 3G). Interestingly, increasing values of these terms showed the opposite relationship with response times in the second response state after a rare transition. In other words, the higher the inferred chosen option probability (which assumes a common transition) the slower the response time when that planned option was interrupted with a rare transition. These findings provide independent behavioral evidence consistent with the MB-HRL framework and the use of options that assume a common transition in this task context.

### The brain represents necessary subgoal, goal, and choice signals for MB-HRL

The MB-HRL framework requires processing subgoals as reinforcing outcomes that signify critical steps towards the goal. To substantiate the neural implementation of the cognitive mechanisms proposed by the model, we first looked for regions of the brain that responded to the completion of subgoals, goals, or both. We designed a whole-brain general linear model (GLM) that included subgoal as a parameteric modulator locked to onset of the outcome state and a regressor for the goal presentation at the completion of a fare. At the group level, whole-brain analyses with cluster-level family-wise error (FWE) correction revealed clusters positively correlated with subgoal collection most notably in basal ganglia (caudate, putamen and nucleus accumbens), posterior cingulate cortex (pCC), and rACC, extending into vmPFC (Figure 4A, Table 1). There were also negatively correlated clusters in bilateral dmFC (Figure S4A). For goal collection, we found positively correlated clusters along inferior frontal gyrus, in pCC, and in rACC/vmPFC (Figure 4B). A conjunction analysis of the subgoal and goal collection regressors revealed a common activation in rACC/vmPFC and pCC (Figure 4C). This result may suggest some regional overlap (at the level that can be assessed with fMRI) in processing of internally designated subgoals and external reinforcers. We also programmed a parametric modulator for proximity to goal, in terms of subgoals, that was time-locked to both of the response states. There were positive correlations with this indicator of abstract progress towards goal in bilateral insula and right frontal pole (Figure 4D). Although, this representation may not be entirely linear, particularly in frontal pole. These signals provide critical representations for subgoal processing in MB-HRL processes.

**Table 1.** Neural signatures of MB-HRL. Related to Figures 4 and 6, a table of the GLM results with relative exploration-weighted value as the decision variable. We used a cluster-forming threshold of *p <* 0.001 uncorrected, followed by a minimum cluster size instantiating a whole-brain cluster-level, FWE-corrected threshold of *p <* 0.05. Red indicates a positive correlation with the regressor and blue a negative correlation.

| peak region | hemisphere | x | y | z | p(FWE-corr)<br>cluster-level | cluster size<br>(voxels) | T | Z |
| --- | --- | --- | --- | --- | --- | --- | --- | --- |
| **current subgoal** |  |  |  |  |  |  |  |  |
| insula | R | 30 | 26 | -5 | 5.4978E-07 | 735 | 10.35 | 6.59 |
| insula | L | -31 | 24 | -3 | 7.9091E-05 | 442 | 7.83 | 5.65 |
| frontal pole | R | 28 | 60 | -9 | 6.5336E-08 | 875 | 7.52 | 5.52 |
| lateral occipital cortex | L | -33 | -63 | 62 | 7.0297E-09 | 1030 | 6.8 | 5.18 |
| caudate | R | 10 | 8 | 18 | 0.00036681 | 362 | 5.87 | 4.7 |
| superior frontal gyrus | R | 2 | 26 | 56 | 4.5575E-07 | 747 | 5.35 | 4.4 |
| middle frontal gyrus | R | 48 | 26 | 44 | 0.0006311 | 335 | 5.21 | 4.32 |
| precuneous cortex | R | 4 | -57 | 46 | 0.03411664 | 159 | 5.1 | 4.26 |
| lateral occipital cortex | R | 12 | -73 | 46 | 0.0430529 | 150 | 4.59 | 3.93 |
| vmPFC | R | 2 | 32 | -13 | 1.4111E-05 | 538 | -6.88 | -5.22 |
| occipital fusiform cortex | L | -25 | -71 | -15 | 0.00117344 | 305 | -5.74 | -4.63 |
| temporal occipital fusiform cortex | R | 28 | -57 | -19 | 5.2371E-08 | 890 | -5.59 | -4.54 |
| central opercular cortex | L | -59 | -23 | 16 | 0.00015578 | 406 | -5.14 | -4.28 |
| supramarginal gyrus | R | 52 | -25 | 28 | 0.00555416 | 234 | -4.99 | -4.19 |
| lingual gyrus | R | 4 | -77 | -9 | 0.01459937 | 193 | -4.77 | -4.04 |
| **goal amount** |  |  |  |  |  |  |  |  |
| inferior temporal gyrus | L | -65 | -27 | -27 | 0.00298554 | 234 | -4.79 | -4.06 |
| **relative exploration-weighted value** |  |  |  |  |  |  |  |  |
| rACC | R | 4 | 44 | 10 | 8.6931E-06 | 480 | 5.54 | 4.52 |
| MTG | L | -69 | -51 | 8 | 0.00057481 | 290 | 5.28 | 4.36 |
| superior temporal gyrus | R | 60 | -33 | 4 | 0.00082811 | 275 | 5.09 | 4.25 |
| superior parietal lobule | R | 16 | -49 | 76 | 0.0057393 | 200 | 4.69 | 3.99 |
| vIPFC | R | 42 | 38 | -7 | 0.04778206 | 127 | 4.46 | 3.85 |
| dmFC | R | 4 | 16 | 54 | 6.5306E-12 | 1317 | -7.13 | -5.34 |
| superior frontal gyrus | R | 26 | 2 | 56 | 0.00016212 | 344 | -6.02 | -4.78 |
| frontal operculum cortex | L | -35 | 26 | 6 | 0.0094394 | 182 | -5.73 | -4.62 |
| lateral occipital cortex | L | -13 | -77 | 56 | 0.00012632 | 355 | -5.34 | -4.4 |
| precuneous cortex | R | 22 | -63 | 24 | 0.01365019 | 169 | -4.79 | -4.06 |
| **rare transition > common transition** |  |  |  |  |  |  |  |  |
| temporal occipital fusiform cortex | R | 30 | -55 | -11 | 6.7848E-10 | 1038 | 7.4 | 5.47 |
| temporal occipital fusiform cortex | L | -33 | -49 | -21 | 7.9936E-15 | 1851 | 7.09 | 5.32 |
| lateral occipital cortex | R | 38 | -83 | 22 | 0.0053824 | 207 | 4.72 | 4.02 |
| MTG | R | 64 | -39 | 8 | 0.00281263 | 232 | -6.2 | -4.88 |
| supramarginal gyrus | R | 52 | -39 | 30 | 0.00150301 | 257 | -5.48 | -4.48 |
| **subgoal collection > no subgoal collection** |  |  |  |  |  |  |  |  |
| accumbens | R | 12 | 8 | -13 | 2.2204E-16 | 2307 | 9.67 | 6.36 |
| rACC | L | -3 | 40 | 2 | 4.0412E-14 | 1864 | 8.83 | 6.06 |
| lateral occipital cortex | L | -37 | -89 | -9 | 8.3549E-08 | 801 | 7.68 | 5.59 |
| lateral occipital cortex | R | 40 | -83 | -11 | 2.8238E-11 | 1349 | 7.42 | 5.47 |
| pACC | R | 2 | -25 | 30 | 4.4367E-07 | 699 | 7.15 | 5.35 |
| superior parietal lobule | L | -43 | -43 | 62 | 0.00367619 | 237 | 5.65 | 4.58 |
| supramarginal gyrus | L | -53 | -25 | 36 | 0.00282932 | 248 | 5.49 | 4.48 |
| frontal pole | L | -17 | 66 | -15 | 0.04032811 | 144 | 4.25 | 3.7 |
| intracalcarine cortex | R | 14 | -81 | 8 | 0 | 6451 | -10.85 | -6.74 |
| superior frontal gyrus | R | 10 | 4 | 66 | 1.3188E-12 | 1583 | -8.09 | -5.76 |
| frontal pole | L | -41 | 46 | 22 | 0.00891412 | 201 | -6.69 | -5.13 |
| insula | R | 32 | 20 | 8 | 0.03146645 | 153 | -6.65 | -5.11 |
| frontal operculum cortex | L | -43 | 12 | 6 | 0.00022224 | 363 | -6.21 | -4.88 |
| **SPE** |  |  |  |  |  |  |  |  |
| intraparietal sulcus | L | -33 | -53 | 44 | 0 | 7241 | 9.4 | 6.27 |
| precentral gyrus | L | -43 | -3 | 40 | 0 | 2401 | 8.4 | 5.89 |
| inferior temporal gyrus | R | 48 | -55 | -15 | 2.2204E-16 | 2255 | 8.08 | 5.76 |
| inferior temporal gyrus | L | -63 | -55 | -13 | 1.9984E-15 | 2059 | 6.63 | 5.1 |
| superior frontal gyrus | R | 24 | 16 | 52 | 2.002E-06 | 595 | 5.8 | 4.66 |
| precentral gyrus | R | 40 | 8 | 30 | 0.00595602 | 212 | 5.62 | 4.56 |
| parietal operculum | L | -59 | -29 | 18 | 2.336E-11 | 1326 | -7.95 | -5.7 |
| cerebellum | L | -29 | -83 | -33 | 0.00029969 | 340 | -7.94 | -5.7 |
| insula | R | 40 | 10 | -5 | 1.3666E-07 | 750 | -7.27 | -5.4 |
| superior parietal lobule | L | -25 | -49 | 70 | 0 | 2624 | -6.98 | -5.27 |
| superior parietal lobule | R | 20 | -45 | 74 | 0.00276205 | 243 | -6.51 | -5.04 |
| supramarginal gyrus | R | 68 | -29 | 30 | 2.0828E-12 | 1505 | -6.24 | -4.9 |
| frontal pole | L | -9 | 66 | 28 | 1.0132E-11 | 1387 | -5.88 | -4.71 |
| postcentral gyrus | L | -61 | -17 | 46 | 0.04246481 | 139 | -5.51 | -4.5 |
| cerebellum | R | 32 | -83 | -31 | 0.00188117 | 259 | -5.12 | -4.26 |
| goal collection |  |  |  |  |  |  |  |  |
| lingual gyrus | L | -1 | -89 | -11 | 0 | 14658 | 16.9 | inf |
| inferior frontal gyrus | R | 46 | 10 | 28 | 1.8763E-14 | 1983 | 8.47 | 5.92 |
| frontal orbital cortex | R | 24 | 30 | -17 | 4.3771E-08 | 865 | 7.64 | 5.57 |
| pACC | R | 8 | -49 | 34 | 0.00431935 | 236 | 6.11 | 4.83 |
| pACC | L | -11 | -45 | 36 | 0.03458422 | 153 | 6.04 | 4.79 |
| lateral occipital cortex | R | 10 | -63 | 62 | 0 | 16229 | -15.38 | -inf |
| precuneous cortex | L | -17 | -63 | 14 | 0 | 4689 | -11.18 | -6.84 |
| cerebellum | R | 36 | -43 | -39 | 3.3211E-12 | 1554 | -7.72 | -5.61 |
| lateral occipital cortex | R | 44 | -79 | 38 | 0.04064624 | 147 | -7.2 | -5.37 |
| frontal operculum cortex | L | -35 | 24 | 10 | 4.3944E-06 | 582 | -6.75 | -5.16 |
| caudate | L | -5 | 4 | 14 | 0.00028063 | 361 | -6.48 | -5.02 |
| temporal pole | L | -51 | 14 | -25 | 6.9868E-05 | 431 | -6.05 | -4.8 |
| frontal pole | R | 32 | 44 | 30 | 0.00509717 | 229 | -5.89 | -4.71 |
| brainstem | L | -9 | -27 | -45 | 0.00534568 | 227 | -5.87 | -4.7 |
| cerebellum | L | -37 | -47 | -41 | 0.00846039 | 208 | -5.43 | -4.45 |
| temporal pole | R | 36 | 16 | -33 | 0.0039322 | 240 | -4.59 | -3.93 |

**Figure 4.**
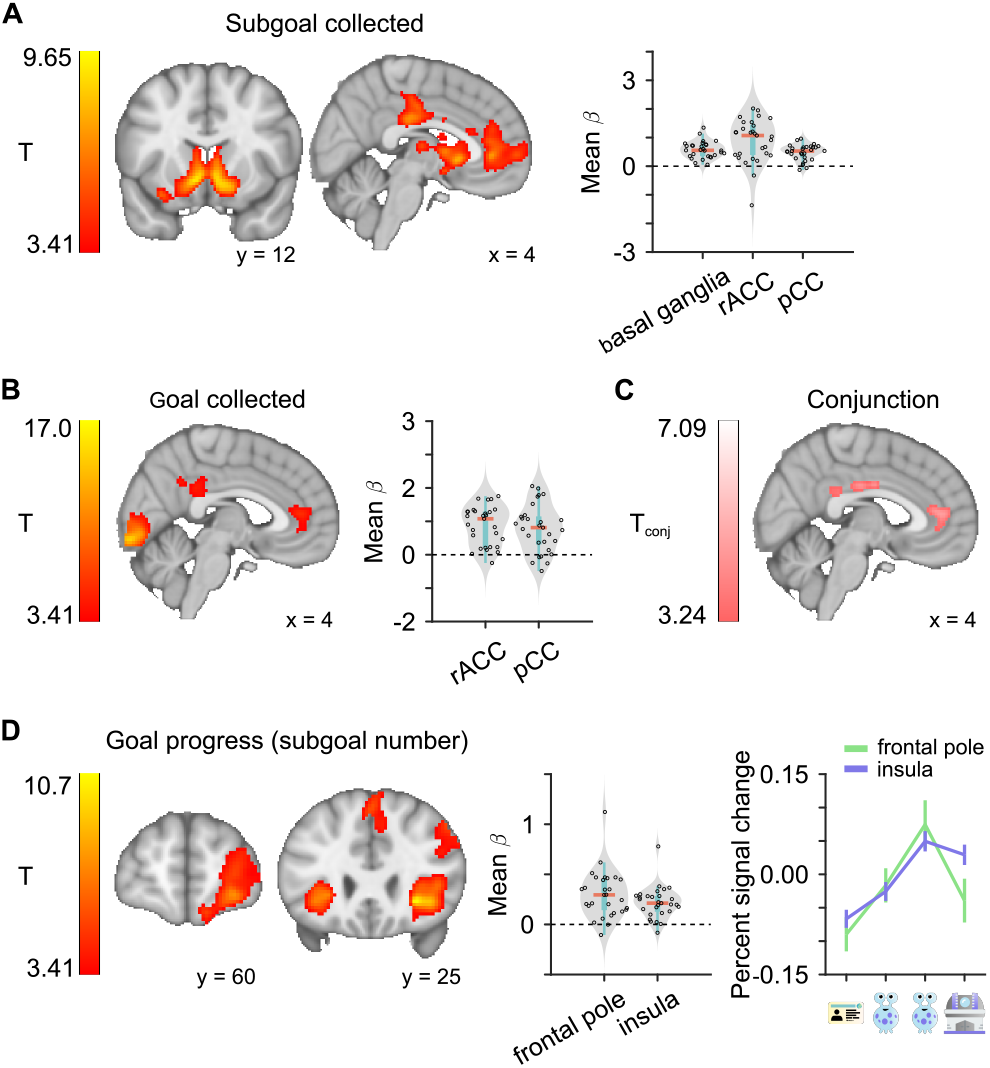
Encoded neural signatures of MB-HRL. (A) Positive correlation with subgoal collection from the GLM, using a cluster-forming threshold of *p <* 0.001 uncorrected, followed by cluster-level FWE correction at *p <* 0.05. Left: population-level T-statistic maps. Right: Post-hoc plots of individual-level regression coefficients extracted from within the significant clusters defined at the population level to demonstrate variance in the reported effects across participants. Top and bottom edges of blue boxes represent the 75th and 25th percentile, respectively. Orange line is the median of the data. (B) Positive correlation with goal collection, as in A. (C) Map of significant T-statistics for conjuntion of subgoal and goal collection. (D) Left, middle: Positive correlation for goal progress, as in A, B. Right: Mean, within-cluster percent signal change across subjects for each subgoal to demonstrate the response profile of the goal progress effect). Error bars represent SEM.

Representing the current subgoal is crucial to computing values appropriate to the task sequential context. To examine latent representations of the current subgoal, we sought to decode this information during the first response state, when the option would be initiated (Figure 5A). We performed a searchlight multivoxel pattern analysis (MVPA), training a linear decoder (support vector classifier) on neural activity in spheres of voxels centered on every voxel in the brain. The decoder was trained on activity during the outcome state, with the label of the *encountered* subgoal. Essentially, we looked for representations of the current subgoal during decision-making that were similar to how that subgoal was represented when it was experienced. This training procedure may miss some representations of current subgoal that are distinct from experiential representations, but it circumvents the confounding correlation between current subgoal and goal progress. Using a very stringent test for significance at the population level, we tested for significant prevalence of above-chance decoding accuracy across subjects using a minimum statistic approach (see Methods; Allefeld et al., 2016). We found significant clusters in the right cuneus, right vmPFC, and right insula/central operculum (cO; Figure 5B,C). In a significant majority of these subjects, these regions contain information about the current subgoal during initiation of the option that is represented in the same fashion as when that subgoal is encountered. This information about latent task structure could be used to orient choices towards the correct subgoal in enabling hierarchical behavior.

**Figure 5.**
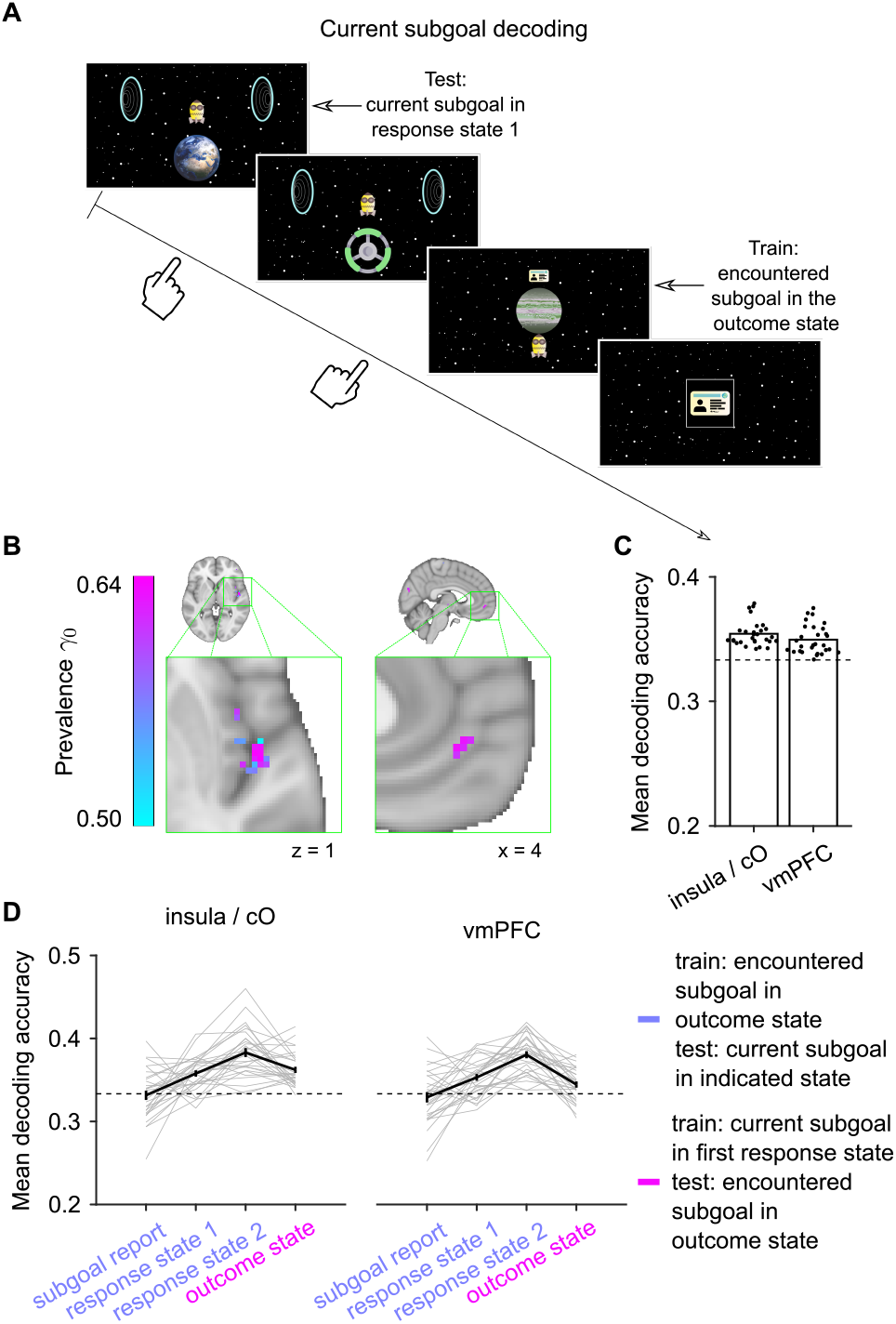
Decoding latent subgoal representations during hierarchical behavior. (A) Schematic of the approach to whole-brain searchlight decoding of current subgoal during the first response state. (B) Clusters of significant prevalence of current subgoal decoding from the whole-brain searchlight analysis. Color indicates the probability at which the prevalence null can be rejected. (C) Mean individual-level decoding accuracies within the significant prevalence clusters (black dots). Bars represent group mean and the dashed line is theoretical chance (*P* = 1*/*3). (D) ROI decoding accuracies across task epochs within spheres centered on the significant prevalence results for individuals (gray lines) and group average (black line, error bars are SEM). Dashed line is theoretical chance (*P* = 1*/*3). Note that accuracies from different decoding schemes (indicated by purple and pink labels) may not be directly comparable.

To better understand how the current subgoal was represented over the course of each trial, we ran another decoding analysis within the insula/cO and vmPFC regions we identified from the searchlight analysis. We used spherical ROIs centered on voxels with the highest probability at which the prevalence null could be rejected and decoded the current subgoal in these regions during different task epochs. We found that subgoal decoding accuracies were robust throughout the trial in both regions, but not during the subgoal report (Figure 5d). We note, however, that because the subgoal report only happened only on 1*/*3 of trials, the decoding results for that particular event may have been underpowered relative to the other trial components. These results show that the latent representation of current subgoal persists across decision behavior and outcomes. With little and indirect feedback about what is the current subgoal, the maintenance of this representation across time is necessary to enable ongoing selection of behavior towards the appropriate subgoal.

In addition to hierarchy at the level of task structure, evidence from response times suggested that behavior was also hierarchical with regard to actions. We examined neural activity for further evidence that subjects were executing options that assumed a common transition in between actions. We took a similar searchlight decoding approach to test for the presence of information about the second action during the first response state, when the first action was being executed and prior to the state transition (Figure 6A). We trained the linear decoder on neural activity during this state and tested it on held-out data from the same epoch. We found significant prevalence of decoding accuracy in left motor cortex and right amygdala (Figure 6B,C). The consistent presence of this information reveals a plan based on a common transition in areas downstream of decision-making regions. This result also further validates the implementation of options in the MB-HRL model.

**Figure 6.**
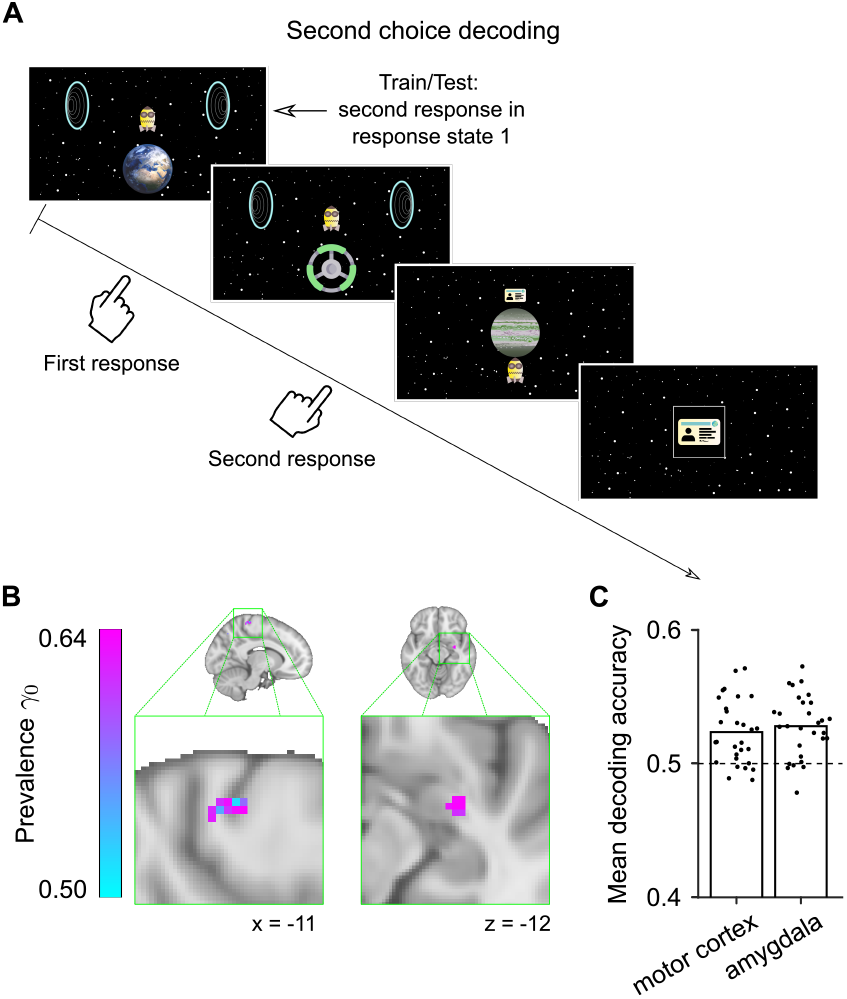
Decoding action plans that rely on knowledge of task structure. (A) Schematic of the approach to whole-brain searchlight decoding of second action during the performance of the first. (B) Clusters of significant prevalence of second action decoding from the whole-brain searchlight analysis. Color indicates the probability at which the prevalence null can be rejected. (C) Mean individual-level decoding accuracies within the significant prevalence clusters (black dots). Bars represent group mean and the dashed line is theoretical chance (*P* = 1*/*2).

### Activity in regions of mFC tracks decision variables during model-based, hierarchical behavior

If the brain uses cognitive mechanisms akin to those proposed by MB-HRL in order to produce the observed behavior, we should expect to see neural correlates of the learning and decision variables generated by the model. To test for these relationships, the GLM described above included two of these variables. Identical to previous model-based formalizations, the MB-HRL agent learns the state-action transition probabilities by computing the difference between its expectation for the resulting state and the actual outcome. This state prediction error (SPE) is then used to update the expectation of the state-action transition probability. SPEs were programmed in the GLM as parametric modulators aligned to the outcome state event regressor. We found positive correlations between neural activity and SPEs most robustly in the intraparietal sulcus and left lateral prefrontal cortex, consistent with previous reports (Figure S6A; Gläscher et al., 2010; Lee et al., 2014; Kim et al., 2019).

We also tested for encoding of decision variables during both response states. Because the decision variables were partially correlated, we ran separate GLMs to test for various forms of value signals: the relative exploration-weighted value of the chosen option (i.e., choice probability, Table 1), exploration-weighted value of the chosen option (Table 2), value of the chosen option, and the relative value of the chosen option. We found significant, cluster-corrected neural correlates of these variables (Figure 7, Figure S6). The clusters for different decision variables were overlapping in some regions, so we performed Bayesian model selection (BMS) to determine which variable explained the most variance in neural activity in these regions. Specifically, we first created ROI masks for each decision variable that survived correction within each ROI. To protect against the possible effects of non-independence bias in the ROI selection, for each region we tested for a given variable not only in the ROI defined by that specific variable, but also in the ROIs defined by other decision variables yielding significant clusters in that specific region. In this way we tested whether one variable best explained variance, even in ROIs defined by other variables. We found that the robust positive correlations in rACC (extending into the paracingulate gyrus; Figure 7A,B) as well as the right ventrolateral prefrontal cortex (vlPFC; Figure S6F,G) were best explained by relative exploration-weighted value irrespective of which variable used to define the ROIs. Interestingly, the rACC cluster substantially overlapped with the regions that responded to subgoals (Figure S6B) and goals (Figure S6C), suggesting a possible updating of these decision signals by the outcomes. The BMS analysis also determined that activity in a positively correlated cluster in middle tempral gyrus (MTG; Figure 6F,G) and in a negatively correlated cluster in dmFC (Figure 7A,B) could best be attributed to chosen exploration-weighted value across all ROIs. These results demonstrate that this activity relates to specific exploration-weighted value signals in each region that incorporate both exploration biases and computed values.

**Table 2.** GLM with chosen exploration-weighted value. A table of the GLM results as in Table 1, but with chosen exploration-weighted value as the decision variable. Results for the other variables are comparable to those in Table 1.

| peak region | hemisphere | x | y | z | p(FWE-corr)<br>cluster-level | cluster size<br>(voxels) | T | Z |
| --- | --- | --- | --- | --- | --- | --- | --- | --- |
| **chosen exploration-weighted value** |  |  |  |  |  |  |  |  |
| MTG | R | 64 | -31 | 2 | 0.0050934 | 209 | 5.29 | 4.37 |
| MTG | L | -69 | -49 | 12 | 0.02257588 | 155 | 4.35 | 3.77 |
| dMFC | R | 4 | 18 | 52 | 1.3179E-05 | 471 | -6.88 | -5.22 |
| middle frontal gyrus | L | -39 | -1 | 64 | 6.3812E-05 | 395 | -6.19 | -4.87 |
| frontal operculum cortex | L | -33 | 26 | 4 | 0.02392085 | 153 | -5 | -4.19 |
| inferior frontal gyrus | L | -57 | 18 | 30 | 0.01900173 | 161 | -4.67 | -3.98 |
| superior frontal gyrus | R | 26 | -1 | 56 | 0.04449345 | 132 | -4.59 | -3.93 |

**Figure 7.**
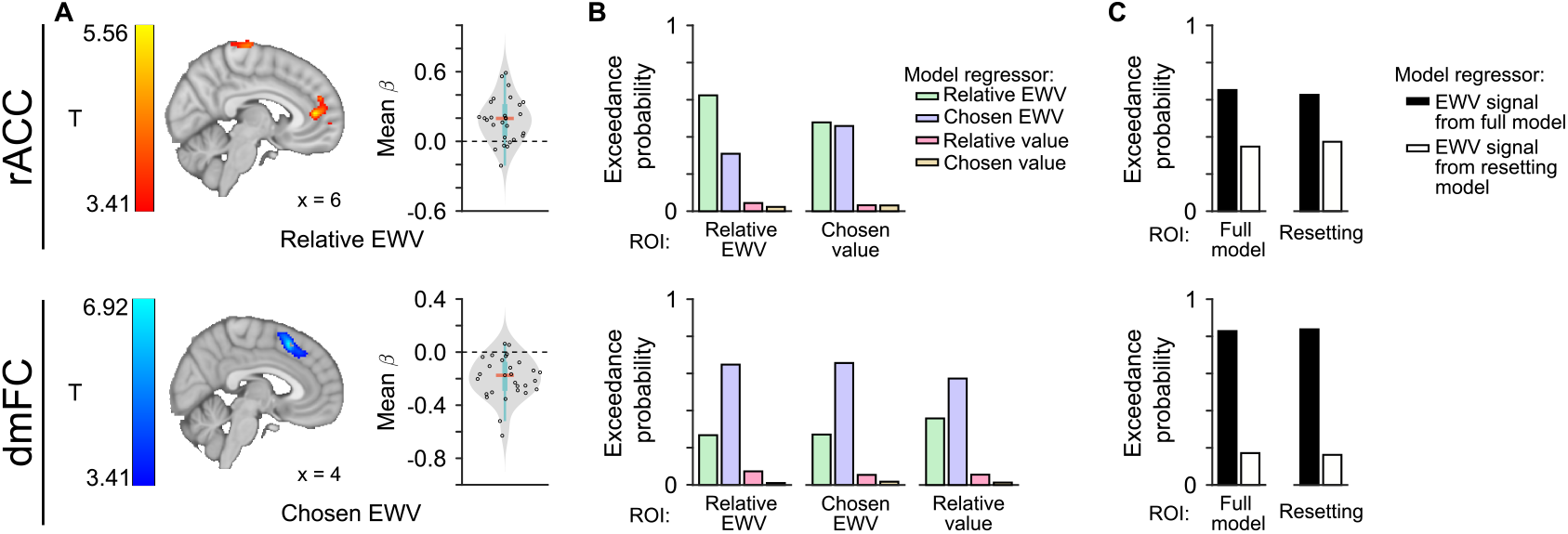
MB-HRL learning and decision variables are represented in neural activity. (A) GLM correlations with decision variables tested using a cluster-forming threshold of *p <* 0.001 uncorrected, followed by cluster-level FWE correction at *p <* 0.05. Left: population-level T-statistic maps. Right: individual-level regression coefficients within the significant clusters defined at the population level. Top and bottom edges of blue boxes represent the 75th and 25th percentile, respectively. Orange line is the median of the data. Top row: positive correlations between rACC and relative exploration-weighted value (EWV). Bottom row: negative correlations between dmFC and chosen exploration-weighted value. (B) Results from a BMS comparison of GLMs to test different value-related variables. Each bar plot shows mean exceedance probabilities for the GLMs with different decision variables in an ROI defined by a significant cluster from one of the GLMs. ROIs were only assessed for variables that survived cluster corrections. ROIs were limited to rACC (top row) and dmFC (bottom row) regions. (C) Results from a BMS comparison of GLMs with (full model) and without (resetting model) latent learning. ROIs were generated in the same fashion as in B.

An alternative interpretation to the negative correlation with chosen exploration-weighted value in dmFC is that this region has a positive correlation with hierarchical planning complexity relative to the goal (Balaguer et al., 2016). In this task, such complexity would decrease across the sequence of subgoals, as fewer steps remain towards the goal. However, the chosen exploration-weighted value decision variable is negatively correlated with the current subgoal, so related dmFC activity decreases over goal progress (Figure S6D). Indeed, activity in the dmFC region has a positive relationship with goal progress (Figure S6E) that has been accounted for in the same GLM.

To provide further evidence that these relative exploration-weighted value signals incorporated model-based values, we ran separate GLMs using exploration-weighted value variables from the resetting version of the MB-HRL model. This version of the behavior model learns the state-action space within a subgoal and contains the same exploration biases as the full version of the model, so there is some overlap in exploration-weighted value signals with the full version. However, the resetting model does not carry its knowledge of the environment into a new subgoal (no latent learning). A similar BMS approach as before determined that the exploration-weighted value signals from the intact model best explained neural activity in dmFC, mPFC, MTG, and vlPFC across ROIs determined by GLMs with value signals from either behavior model (Figure 7C, Figure 6H). Therefore, persistent knowledge of the state-action space—a fundamental feature of model-based decision-making—is important for explaining the exploration-weighted value signals observed in all regions. Thus, we extend the known functionality of these regions of frontal cortex, establishing a relationship between neural activity in these regions and exploration-weighted value of choice during model-based, hierarchical decision-making. Specifically, we show that these regions represent integrated model-based values computed as a function of subgoals, not external reinforcers.

## Discussion

Human decision-making is both model-based and hierarchical. The MB-HRL framework adapted here provides a computational account of how these two aspects of cognition can be integrated (Botvinick and Weinstein, 2014). This model explained choices and response times in a novel, sequential-subgoal decision-making task, capturing several features of model-based, hierarchical behavior. Substantiating this framework, we found signals related to the encoding and processing of subgoals in sequence across several brain regions associated with processing of external reinforcers. Sequential, hierarchical behavior requires a representation of the current subgoal to appropriately drive behavior in the correct order. We decoded a latent representation of the current subgoal from insula/cO and vmPFC and found that it persisted throughout the trial while participants actively perform behavior and receive outcomes. In MB-HRL, values are computed as a function of the subgoal (hierarchical) as well as learned task structure (model-based). Neural activity in the mFC and vlPFC tracked relative exploration-weighted value of the chosen option or action during execution of that choice and activity in another region of mFC tracked the exploration-weighted value of the chosen option. These findings provide evidence for an integration of model-based and hierarchical processes in the human brain providing support for the MB-HRL framework while revealing regions critical for the representation of subgoals and values computed as a function of them.

Subgoals are internally-designated states of the world, distinct from external reinforcers, that can be used as interim goals to reinforce and guide decisions. Previous studies have shown that the experience of external reinforcers results in feedback signals in basal ganglia (O’Doherty et al., 2004; Niv, 2009). Here, we show that activity in basal ganglia (in all three parts of the striatum: the caudate, putamen and nucleus accumbens) responded to the collection of subgoals, potentially consistent with a feedback signal for hierarchical processes Botvinick et al. (2009); Rasmussen et al. (2017); Janssen et al. (2022). We observed that responses to both subgoals and goals encompassed parts of rACC and vmPFC, as well as pCC. These responses may also constitute crucial feedback about outcomes that update the subsequent decision signals. In our task, subgoal and goal collection are correlated with a change in current subgoal, so it may be the case that these signals are also related to a change in planning or strategy. In fact, pCC has been been implicated in integrating outcomes to modulate strategy (Pearson et al., 2009), among other cognitive functions (Foster et al., 2023). The outcome signals in the frontal cortex regions, on the other hand, are in line with a rich literature of goal valuation and decisions over goals in vmPFC (O’Doherty, 2011).

We also decoded representations of subgoal identity from vmPFC, cuneus, and insula/CO. An important feature of this subgoal representation is that it relied on correct knowledge of the task structure, which is necessary to guide decisions in the correct order. The representation was also maintained persistently, over the course of the trial, which may be vital for keeping ongoing behavior aligned to the appropriate subgoal. A previous study reported goal category encoding in the vmPFC during a willingness-to-pay task (McNamee et al., 2013). Together, these results suggest the possibility that in some regions, neural architectures can flexibly incorporate goals and subgoals, depending on task demands, in order to make decisions. Current subgoal was also decoded from activity in precuneus, a region that has been associated with visuo-spatial working memory (Courtney et al., 1996, 2007). Finally, we also decoded a latent representation of the current subgoal in the insula and cO—crucial nodes in the cingulo-opercular network (sometimes called “action-mode network”; Dosenbach et al., 2025) thought to be involved in task control across different conditions (Dosenbach et al., 2006, 2007, 2008). Of particular relevance, some of these studies have demonstrated persistent activity across the task epoch in this network, possibly suggesting the maintenance of a representation of the task set that appropriately guides behavior according to the learned structure of the current task (Dosenbach et al., 2007). Here, the sustained representation of the current subgoal in insula and cO may reflect a task set, driving behavior in the appropriate sequence.

Representing subgoals is necessary for evaluating choices in pursuit of those subgoals. Here, we identified signatures of exploration-weighted value in key frontal regions. The vlPFC tracked the relative exploration-weighted value of the chosen option, which establishes a new function that is consistent with the broader role of lateral prefrontal cortex in cognition and value-based decision-making (for review see Dixon and Christoff, 2014). The more ventral region where we observed this signal has been associated with tracking the reliability of different decision-making strategies in a similar two-step task (Lee et al., 2014; Kim et al., 2019). In that task, the reliability of model-based learning was manipulated by varying the state transition statistics. While the Space Taxi task lacks such manipulations, policy reliability may derive from exploration-weighted value in this task (i.e., the policy is reliable when either the exploration biases or the learned values drive up the exploration-weighted value).

A strong negative correlation with relative exploration-weighted value was found in dmFC, mostly in pre-supplementary motor area, that complements previous reports. Previous neuroimaging during a two-armed bandit (Wunderlich et al., 2009) and two-step task (Lee et al., 2014), for example, demonstrated a negative correlation with the relative value of the chosen stimulus or action in this region. In both of these cases the signal was interpreted as a positive correlation with the relative value of the *unchosen* stimulus or choice. Human single-neuron recordings in this region during a two-armed bandit task showed that individual units tracked the relative value of one stimulus over the other, irrespective of what choice was made (Aquino et al., 2023). The discrepancies between methods of observation signify the importance of more fine-grained observations in clarifying the precise nature of these cognitive representations. The same study also demonstrated that many of these neurons encoded an exploration-weighted value signal that integrated both value and uncertainty. Here, dmFC activity is best explained by chosen, not relative, exploration-weighted value, but future studies will be important for discerning these representations with more granularity.

Activity in the rACC robustly signaled relative exploration-weighted value. This region overlaps with value-related vmPFC activity seen in many studies (Clithero and Rangel, 2013), but extends more dorsally. This region also overlaps with some areas found to have unique involvement in model-based processes across studies (Huang et al., 2020). A few relevant studies have found choice-value-related signals precisely in this region. In a forced choice task with changing response contingencies, rACC was found to track the reliability of an exploitation choice relative to an exploration choice (Domenech et al., 2020). Consistently, the implementation of reliability in their model could also correspond to high relative value. In another study, activity in rACC tracked the immediate values of different choices as well as the value of those choices computed using a planning mechanism that took into account how those choices affected future states of the task (Kolling et al., 2018). A separate line of research has suggested a role for rACC, potentially related to planning, in representing abstract structure in the world, such as “schema” in narrative events (Gilboa and Marlatte, 2017; Baldassano et al., 2018) and “cognitive maps” of conceptual relationships (Constantinescu et al., 2016; Park et al., 2021). One study linked rACC activity to the relative value of two choices computed as a function of a representational map (Park et al., 2021). Our results unify and add key components to these functions, suggesting that rACC integrates an internal model of the world, planned actions or options, and the relative exploration-weighted value of those plans computed as a function of exploration heuristics and subgoals.

Varying theories from Holroyd and colleagues have proffered that anterior cingulate cortex is involved in several aspects of hierarchical behavior. Some of these ideas have situated rACC as representing the more abstract task and action representations in a rostral to caudal hierarchy. They have suggested that rACC may represent hierarchical models of action and goal structure (Holroyd and Verguts, 2021) or selection and maintenance of high-level options or goals (Alejandro and Holroyd, 2024). The human literature that motivated these theories only provide vague and indirect support. Here, we provide empirical evidence in support of this theoretical framework, while also formalizing its implementation and providing a quantitative explanation of rACC function.

In our MB-HRL framework, first-response decisions are modeled as choices over options, though MB-HRL first requires an action model to learn an options model (Sutton et al., 1999; Botvinick and Weinstein, 2014). The response time and choice decoding results support the inclusion of options, but the brain may still simultaneously track action values. In this task, action and option values would be highly correlated, so conclusions about whether the relative exploration-weighted value signal we observe is related to actions or options should be tempered. Prior work has shown (goal-dependent) action value tracking in dmFC (Wunderlich et al., 2009; Lee et al., 2014; Aquino et al., 2023) and vmPFC (Clithero and Rangel, 2013). That being said, what experimentalists typically refer to as single actions are actually composite motor behaviors (e.g., a button press involves a coordinated, complex sequence of muscle fiber activations). Indeed, studies in humans (Roland et al., 1980; Amador and Fried, 2004; Cona and Semenza, 2017) and non-human primates (Tanji and Shima, 1994; Tanji, 2001) have characterized sequential motor representations in pre-supplementary motor area. Thus, future work will need to tease apart value signals related to actions and options, which may require observations of neural activity with greater spatial and temporal precision.

While the neural activity may not resolve action and option values, the behavior and decoding of planned actions evidences the use of options in stochastic environments. Action sequences or options typically involve very high probability state-action transition probabilities. For example, the sequence of actions required to reach for a coffee mug is unlikely to be interrupted by a change in the world. Our MB-HRL model formalization aligns with this intuition, suggesting options form based on statistical regularities in the environment. Indeed, learning of statistical, spatiotemporal regularities in the environment (Fiser and Aslin, 2002) or latent graph structures (Schapiro et al., 2013; Kahn et al., 2018) has been observed previously, and has been shown to influence response times (Hunt and Aslin, 2001; Lynn et al., 2020). Analogously in our task, response time differed between rare and common transitions. We also find that that response times varied and as a function of option value and outcome in a manner that suggests planning on the assumption of a common transition. We also demonstrated an ability to decode the second action during performance of the first. This information was found in motor cortex, a region previously implicated in sequential behavior in many studies (Karni et al., 1998; Tanji, 2001). We also decoded future choice information from the amygdala, which may relate to its known role in tracking states or choices in goal-directed plans (for review see Johnson and Grabenhorst, 2025). In this context, these results demonstrate a novel mechanism by which sequences of choices can be evaluated and planned efficiently by relying on the task structure.

Hierarchical abstraction in state-action spaces aids learning and planning by reducing computational load. In our task, deciding over only common-transition options may help manage working memory demands, as representing the full space is cognitively costly. This demand is heightened by the change in state-action space with each new fare, requiring rapid learning. Task structure may have interacted with working memory constraints to further encourage option use: the stable yet low probability of rare transitions (*P* = 0.3) and forced availability of subgoals on high probability sequences allowed rare outcomes to be ignored without sacrificing performance. Future studies could manipulate environment statistics to better understand option formation in response to those statistics.

Many participants exhibited stereotyped exploration patterns that predicted better task performance. This strategy may ease working-memory load by learning and organizing state-action representations in a consistent order, resulting in improved performance. However, exploration stereotypy could also simply be correlated with some other strategy or cognitive ability that engenders better performance. While exploration biases in MB-HRL resemble novelty bonuses, they are a distinct heuristic. Subjects explored novel states to discover subgoals, but the order followed self-generated patterns rather than just novelty. The consistency across subjects in exploration order (i.e., left-left, left-right, right-left, right-right) suggests that efficient state-action representation may be influenced by learned priors (e.g., reading English from left to right).

Despite MB-HRL’s success in modeling behavior and neural activity, challenges remain. Beyond learning statistical patterns in the environment, a difficult challenge in hierarchical behavior is the discovery and delineation of useful options and subgoals (Sutton et al., 1999; Precup, 2000). Theories posit that subgoals can be learned when they are frequently visited by necessity, constituting bottleneck states (McGovern and Barto, 2001; Stolle and Precup, 2002; Bacon, 2013), as a function of returns (Bacon et al., 2017), or when they enhance planning efficiency (Jinnai et al., 2019; Wan and Sutton, 2022). While some subgoal learning may occur in our task, explicit instructions and practice rounds were designed to mitigate it in the main task that was analyzed. The accurate report of current subgoal and immediate success of many participants in this task evidence the possibility that subgoals can be instructed explicitly, similar to instructed learning (Doll et al., 2009). We relied on these features of the training and behavior to motivate the assumption of correct task structure (i.e., sequence of subgoals) knowledge in our MB-HRL model, leaving aside the difficult problem of option discovery for future studies.

Our MB-HRL model is purely model-based, but other studies have characterized behavior in similar two-step task designs as a mixture of model-based and model-free (Gläscher et al., 2010; Daw et al., 2011; Lee et al., 2014; Kim et al., 2019). Our implementation was motivated by practical considerations. First, latent learning and efficient exploration demonstrate that behavior is largely driven by model-based learning and decision-making. Second, a model-free agent would learn action or option values only as a function of the current subgoal. Consequently, the only instance in which the model-free agent would drive behavior in opposition to the model-based component is in the case of a rare transition on the first trial of the second alien. Such instances are too few upon which a generative model can be conditioned. While performance dipped after these rare transitions, forgetting in MB-HRL was sufficient to capture the difference. Nonetheless, previous studies have characterized behavior with model-free hierarchical reinforcement learning (Botvinick, 2012; Rasmussen et al., 2017; Eckstein and Collins, 2020; Xia and Collins, 2021; Li et al., 2022), so the potential integration of model-free and model-based learning in hierarchical behavior remains an open question.

Another consequence of the MB-HRL framework being purely model-based is an absence of reward and subgoal prediction errors. Previous studies leveraging a model-free hierarchical RL framework demonstrated a positively correlated, unsigned subgoal prediction error signal in dmFC (Ribas-Fernandes et al., 2011, 2019). We observed a negatively correlated, signed response to the outcome in dmFC, however it is hard to compare responses to different variables across such different task conditions. In our case, subgoal collection does not factor in expectations and does not instruct future choices in the same way that subgoal prediction errors immediately influenced behavior in that previous work. But notably, responses to subgoal outcomes occurred in an overlapping dmFC region as the exploration-weighted value signal that we observed at the time of choice, suggesting a possible substrate for the updating of expectations by outcomes. Another study found a hierarchical goal-value error signal in ventral striatum (Diuk et al., 2013); while this signal is not necessarily relevant to our study, we did observe responses to subgoal collection in ventral striatum. Understanding how different learning mechanisms interact in hierarchical behavior remains an important direction for future research.

To conclude, our results provide novel insights into the functions of the frontal cortex, particularly mFC, in hierarchical decision-making involving subgoals. First, we show that the value component of the exploration-weighted value signals reflects latent learning of the state-action structure of the environment. Second, we show that these value signals incorporate heuristic biases in the form of patterned exploration. And third, we show that these decision signals and goal representations are not just related to external reinforcers, but generalize to subgoals: internally-designated states of the world that can be used to reinforce behavior. The results here demonstrate that human behavior integrates model-based and hierarchical processes and that key regions regions evaluate choices as a function of subgoals and model-based knowledge of the environment.

## Acknowledgments

We thank the members of the O’Doherty laboratory for discussions and feedback. We also thank all participants for their participation. This work was supported by the Army Research Office MURI grant W911NF2110328. C.D.G. is supported by the Swartz Foundation Fellowship for postdoctoral research in theoretical neuroscience. J.P.O. is supported by the Tianqiao and Chrissy Chen Center for Social and Decision Neuroscience. The funders had no role in study design, data collection and analysis, decision to publish or preparation of the manuscript.

## Author contributions

C.D.G. collected and analyzed the data. C.D.G., V.M., and J.P.O. designed analyses. C.D.G. and J.P.O. designed experiments and wrote the paper.

## Methods

### Participants

We recruited 37 (18 female) participants from Pasadena and the greater Los Angeles, California area that had previously registered in the Chen Participant Center (between ages 21-66, mean 39 ± 13). All participants had to prove receipt of COVID-19 vaccination and not present any cold, virus, or flu symptoms on the day of participation. Subjects were asked to refrain from recreational drug and alcohol consumption the day prior to and the day of the experiment. Subjects were also encouraged to limit caffeine intake to a typical level on the day of scanning. All participants spoke English, had normal or corrected-to-normal vision, met MRI safety criteria, and reported no history of neurological disorder. Participants were compensated $30 per hour and were given bonuses that equaled the sum of all fares completed ($11 ± 3.7).

### Task

In a room outside of the scanner, subjects first were instructed how to perform the task with instructions on a laptop programmed with the Psychtoolbox (http://psychtoolbox.org/) package for MATLAB. Subjects navigated through the instructions with the keyboard and had to complete a quiz correctly in order to finish the instructions. After, subjects had the opportunity to ask any questions. Subjects were then given an unlimited number of trials to complete 2 practice fares of the game and were given a second chance to ask any more questions. In the fMRI scanner, subjects were run through 3-4 blocks of the task corresponding to 3 blocks during separate fMRI runs and 1 block during collection of structural images, if required. Structural images were not collected for participants that had recently been imaged in another study. In each of the 3 fMRI blocks, participants completed as many fares as they could in 74 trials. In the structural imaging block, subjects had 40 trials. Subjects were given the opportunity to take a break between runs if needed.

At the beginning of each fare in the task, the subjects were shown a screen telling them the value of the next fare ($0.10, $0.30, or $1.00), for 5 s. At the beginning of each trial, a fixation cross was presented for a variable duration, calculated as the sum of the differences between the maximum response times and the actual response times for both states on the previous trial (3.18 ± 0.26 s). On 2*/*3 trials, subjects were then presented with the first response state, signaled by an image of Earth, and given a maximum of 1.8 s to choose between a left and right portal with a corresponding left or right button press. The rocket was animated to move towards the chosen portal with varying duration (1-4 s, randomly sampled from uniform, continuous distribution) in order to jitter the timing of the task epoch. The second response state was then revealed, one of two space stations, contingent upon the choice and probabilistic state transitions. Subjects again had 1.8 s to make a choice and the choice was visualized with the rocket animation of variable duration (1-4 s). The second choice led deterministically to the outcome state. The outcome state with the planet and subgoal (or nothing) was then shown for 1-4 s. The outcome state was followed by a 1 s screen with a white square that contained a subgoal only if it was collected on that trial, but was empty otherwise.

On the other 1*/*3 of trials, participants were first asked to report which subgoal they thought was the current one before being presented with the first response state. The screen showed three random shapes in random position and with different sizes (i.e., small, medium, and large). During training, subjects were instructed to choose the shape with the size that corresponds to the current subgoal, such that small corresponded to permit, medium corresponded to alien, and large corresponded to home. Subjects were informed that their response did not affect the task. Subjects were given 2 seconds to select a shape with a left, right, or middle button press. When errors in this report were made, subjects choices more often aligned with the actual current subgoal than the incorrectly reported one (Figure S1D). This suggests that these errors were a consequence of the reporting mechanism and not incorrect knowledge of the current subgoal.

When the subjects completed a fare, they were shown a goal completion screen for 5 s with the value of the fare acquired. Differences in performance led to differences in task timing, as a result of the goal completion and fare indication screens. As such, some variable down time (197 ± 79 s) was programmed in between runs. During this time, the subjects were shown a screen with the total amount of fare money they had collected so far and the amount of time they had to rest before the next run.

### Data analysis

Analyses were performed with MATLAB (Mathworks) and Python. Data are presented as mean ± standard deviation unless reported otherwise.

### Data analysis: inclusion criteria

To warrant inclusion in the study, subjects had to complete all 3 runs of the task in the fMRI scanner (2 subjects excluded). Subject behavior was also compared to 1, 000 sessions of simulated behavior from an agent that selected actions randomly for all decisions (*P* (*leftchoice*) = 0.5). We calculated the the rare-transition-corrected number of trials per fare (see Data analysis: behavior) for subjects and the corrected trials per fare for all simulated session. The distributions were not normally distributed so we performed a one-sided Wilcoxon rank sum test for each subject compared to the random agent data, to test if their performance was significantly better than chance. Participants were excluded if their performance was not significantly better than chance (6 subjects excluded).

### Data analysis: behavior

As noted in the figure legends, some of the behavior metrics, like the probability of repeating an incorrect sequence (Figure 1E, Figure 3D) and latent learning (Figure 2A, Figure 3B) were limited to common transition trials. These analyses were restricted in this way since there were too few analogous instances with rare transition trials and to avoid effects resulting from a possible strategy of ignoring rare transition trials. For visual purposes, trials to complete a fare over the session (Figure 1F, Figure 3E) were corrected to account for the number of rare transition trials that occurred within that fare. Specifically, the number of rare transition trials without subgoal collection were subtracted from the number of trials in that fare. Since the number of these trials were stochastic and increased the number of trials per fare (effect of rare transition trials: coefficient = 1.64, *t*_760_ = 32.2, *p* = 0), they were removed to get a more reliable sense of actual performance. These effects of rare transition trials and fare number in session were quantified with a linear mixed-effects model with random effects for subjects, given by the following Wilkinson notation:

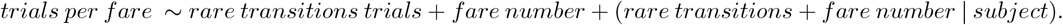

We tested for an effect of goal value on performance using the rare-transition-corrected trials per fare averaged within goal value condition for each subject:

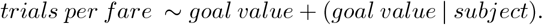

The effects of goal value and trial number on first state response times (RT1) were also asssed by a linear mixed-effects model according to:

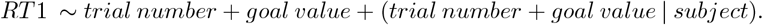

For examining forgetting, the probability of repeating a suboptimal sequence was characterized as a function of how recently that sequence was taken, controlling for the improvement across the session:

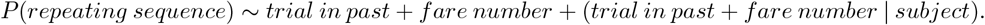

A logistic regression model was designed to examine differences in choice behavior on rare transition trials between first and second alien collection, as well as on the first trial of a new subgoal (when there is potential expression of latent learning). Whether the choice sequence matched the correct one was predicted as a function of trial type (intervening rare transition trial or subsequent common transition trial) while controlling for forgetting (effect of trial number from first alien or first trial of new subgoal):

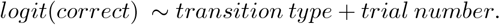

Another linear mixed-effects model was employed to quantify the effects of transition type and trial number on second state response time (RT2):

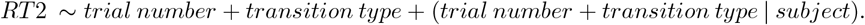

We also quantified the effects of behavior model choice probabilities on response times, accounting for changes over the session and random subject effects according to the linear mixed-effects model:

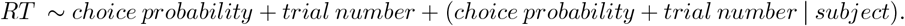

Chosen exploration-weighted value was modeled as a function of current subgoal number in the sequence with another linear mixed-effects model:

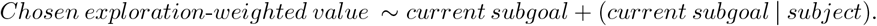

### Data analysis: generative models of behavior

We adapted an MB-HRL model (Botvinick and Weinstein, 2014) for application to behavior in the Space Taxi task and added several parameters to better capture observed behavior. In a model-based fashion, the agent learns and updates a matrix *T* of the probabilities of state-action-state transitions. The transition probabilities for the terminal states are initialized at chance (*P* = 0.25), given that there are 4 possible states (each of the 3 subgoals or nothing). In order to update these expectations with experience, an SPE, *δ*_*SP E*_, is computed as a function of the transition probabilities:

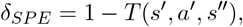

where *s*^*′*^ is the current, second state, *a*^*′*^ is the action taken, and *s*^*′′*^ is the consequent, terminal state. The SPE updates the transition matrix for the given state-action-state probability according to:

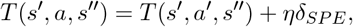

where *η* controls the rate at which the transition matrix is updated. Here, we fix *η* = 1 since learning saturated immediately and then decayed (Figure 1E, Figure S1a). The imperfection in immediate learning and subsequent decay was captured by the forgetting rate *ζ*, a free parameter that draws the transition probabilities back towards chance on every trial:

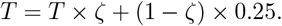

In the first response state, the agent computes the values of options, *o*, as a function of the current subgoal:

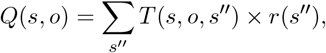

where *s*^*′′*^ is the terminal state and *r* is the pseudo-reward function in which *r*(*s*^*′′*^) = 1 for the current subgoal and 0 otherwise. These options assume the more common path through state-action space, T(*s, a, s*^*′*^, *a*^*′*^, *s*^*′′*^), in which *s*^*′*^ is the higher probability consequence of *a*. This assumption exploits the statistical regularity in the environment to simplify and limit the number of value computations the agent has to make. Previous studies have similarly assumed that subjects know the transition probabilities between the first and second response states as a function of training and experience in the task (Gläscher et al., 2010; Daw et al., 2011; Lee et al., 2014; Kim et al., 2019). Here, we assume that the subject knows which second response states are high probability and which are low. Given this knowledge, if a rare transition occurs, the option terminates according to a deterministic termination function. Values of the two available actions are then similarly computed as a function of the learned structure and the current subgoal:

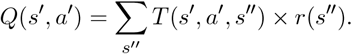

In the first response state, option values are used to generate choice probabilities via a softmax function:

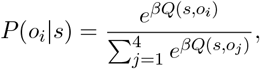

where *β* is the inverse temperature parameter that controls the stochasticity of choice. Here, we fix *β* = 8 due to interactions with the forgetting rate parameter. On rare transitions, the probabilities of actions are also computed according to a softmax function:

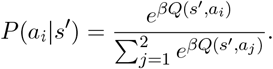

We also found that many subjects expressed stereotyped patterns in exploration behavior. To capture this phenomenon we programmed the agent to compute option utilities, *U*, as a function of values and exploration biases:

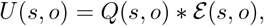

where the exploration biases, E, consist of 4 free parameters that determine the bias towards each of the options until that option has been explored. The option utilities are then fed into the decision function to yield choice probabilities:

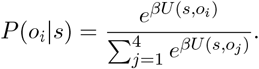

We also observed that the behavior of some subjects improved over the course of the session. The agent was able to capture this effect by defining forgetting rate as a variable that is a function of trial in session:

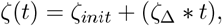

where *ζ*_*init*_ and *ζ*_Δ_ are free parameters that determine the initial forgetting rate and how that rate changes as function of trial number in session, *t*, respectively. The effective forgetting rate was bounded between 0 and 1:

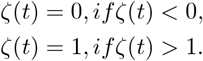

Because we did not observe an effect of fare value on behavior, the MB-HRL model was not programmed to consider goal value. This lack of an effect is likely due to the fact that the fare is not yoked to performance within that fare, and so is independent of how quickly the subjects complete it. Also, to maximize reward across fares, the optimal behavior is to complete as many as possible within the limited number of trials, regardless of the value of any individual fare.

### Data analysis: generative models for comparison

We programmed a few models to demonstrate certain principles of the observed behavior. The random agent was designed to provide a lower bound on performance and to determine the inclusion criterion for performance. This agent selected actions randomly (*P* (*leftchoice*) = 0.5) during both response states on every trial. The optimal agent was designed to provide an upper bound on performance. This agent was the same as the MB-HRL agent characterized above, but had perfect memory (*ζ*= 1), explored randomly, and exploited the highest value option deterministically. The resetting model for Figure 2 and Figure S2 was an MB-HRL agent with perfect memory (*ζ*= 1), explored randomly, and made exploiting decisions (*β* = 8). The resetting model for Figure 3, Figure S3, and Figure 5, was nearly identical to the full MB-HRL agent and whose forgetting and exploration parameters were fit to actual behavior. For both resetting models, with each new subgoal the learned transition matrix was reset to chance (∀(*s*^*′*^, *a*^*′*^, *s*^*′′*^), *T* (*s*^*′*^, *a*^*′*^, *s*^*′′*^) = 0.25).

### Data analysis: model fitting

Models were fit with the probabilistic programming language, Stan (https://mc-stan.org/) via the MatlabStan (https://mc-stan.org/users/interfaces/matlab-stan) interface for MATLAB (Mathworks). We constructed hierarchical models with population-level hyperparameters governing subject-level parameters as a way to enable partial pooling of the data. As a type of regularization, partial pooling helps to avoid overfitting to noise in individual subject sessions without entirely ignoring subject-level variance. The subject-level free parameters that determiend forgetting from the MB-HRL model were modeled as draws from a Gaussian population-level distribution with weakly informative priors (mean *µ* ∼ *N* (0, 1) and variance *σ* ∼ *half* -*Cauchy*(0, 2.5)), under the assumption that subjects behave somewhat similarly over the population. The exploration biases were only modeled at the level of the subject to better capture the heterogeneity in patterns of exploration. The priors on all subject-level parameters were uniform across all possible values, providing no information about what MB-HRL parameter values were likely to be (*ζ*_*init*_ ∈ [0, 1], *ζ*_Δ_ ∈ [−0.01, 0.01], ε_1*−*4_ ∈ [1, 5]). To avoid issues in sampling bounded parameters during fitting, the sampler explored parameters in an unbounded space and transformed those parameters into bounded values by a standard normal inverse cumulative density function. Posterior distributions of parameter estimates were generated in Stan using full Bayesian statistical inference enabled by Hamiltonian Markov chain Monte Carlo sampling (Carpenter et al., 2017). Models were fit using the default no-U-turn sampler, and a Metropolis acceptance rate of 0.9 to force smaller step sizes than the default setting as well as improve sampler efficiency. After 5, 000 warmup draws, models were fit with 10, 000 iterations on 7 parallel chains. For all other settings, default configurations were used.

### Data analysis: estimating parameters, generating variables, and simulating behavior

For comparison of singular parameter estimates to metrics of behavior (Figure S3B), we approximated the mode of subject-level parameter distributions in order to estimate the maximum *a posteriori* parameter values. This approximation was done by binning each parameter distribution in 50 bins and computing the median value of the most populated bin. We generated model variables (e.g., relative exploration-weighted value of the choice) from the model fits at the subject level for comparison to the neuroimaging data. We took all 10, 000 of the Hamiltonian Markov Chain Monte Carlo samples from the subject-level posteriors and used those parameters in an MB-HRL agent exposed to the same task outcomes and forced to make the same choices as the actual subject. The variables were then averaged across all sample runs. We also simulated behavior using these MB-HRL model fits to perform posterior predictive checks. Within each analysis, 1, 000 draws were taken from the subject-level posterior distributions for all parameters, plugged into the relevant model agent, and run through 1 simulation of the task. For the simulation of change in behavior across the session (Figure 3E), the number of fares completed by the agent was matched to the number actually completed by the subject. For all other MB-HRL simulations, the agent was used to simulate behavior for 30*fares*. For random and optimal agent simulations of behavior, those agents were run through 1, 000 iterations of the task with 30*fares*.

### Data analysis: model comparison

In order to compare the predictive accuracy of the MB-HRL models across all choices, we used five-fold cross-validation. For each model, we randomly split each subject’s data into 5 folds. We fit the hierarchical model on 4*/*5 of the folds for all subjects simultaneously, and tested the fit model performance on the left out fold. This procedure was repeated so that all 5 folds were used as the test set. For each fit, after 2, 000 warmup draws, the models were fit with 4, 000 iterations. Otherwise, the fitting procedure was the same as described above. Then, we used parameter samples to calculate log pointwise (per choice) predictive densities for the relevant left out data for each fit. The log pointwise predictive densities were summed across the 5 folds and normalized by number of choices.

### fMRI data acquisition

Neuroimaging data were acquired at the Caltech Brain Imaging Center (Pasadena, CA) using a Siemens Prisma 3T scanner with a 32-channel radiofrequency coil. The whole-brain functional scans were collected with a multi-band echo-planar imaging (EPI) sequence with the following parameters: 72 slices per volume, a repetition time (TR) of 1.12 s, an echo time (TE) of 30 ms, multi-band acceleration of 4, a 54^*°*^ flip angle, a field of view (FoV) of 192 × 192 mm, a 96 × 96 matrix, and 2 mm isotropic resolution. After each run, positive and negative polarity, single-band EPI-based field maps were collected using similar parameters to the functional sequence, but with a TR of 5.13 s, a TE of 41.40 ms, and a 90^*°*^ flip angle. T1- and T2-weighted structural images were collected for all subjects, either at the end of the session or in an earlier session. Both types of structural images were collected with a 230 × 230 mm FoV, a 256 × 256 matrix, and 0.9 mm isotropic resolution. The T1-weighted images were acquired with a TR of 2.55 s, a TE of 1.63 ms, an inversion time of 1.15 s, and an 8^*°*^ flip angle. The T2-weighted images were acquired with a TR of 3.2 s, a TE of 564 ms, and a 120^*°*^ flip angle.

### Data analysis: fMRI data preprocessing

Results included in this manuscript come from preprocessing performed using *fMRIPrep* 22.1.1 (Esteban et al., 2019, RRID:SCR 016216), which is based on *Nipype* 1.8.5 (Gorgolewski et al., 2011, RRID: SCR 002502).

#### Preprocessing of B_*0*_ inhomogeneity mappings

A total of 2 fieldmaps were found available within the input BIDS structure for this particular subject. A B_0_ nonuniformity map (or *fieldmap*) was estimated from the phase-drift map(s) measure with two consecutive GRE (gradient-recalled echo) acquisitions. The corresponding phase-map(s) were phase-unwrapped with `prelude` (FSL 6.0.5.1:57b01774). A B_0_-nonuniformity map (or *fieldmap*) was estimated based on two (or more) echo-planar imaging (EPI) references with `topup` (Smith et al., 2004, FSL 6.0.5.1:57b01774).

#### Anatomical data preprocessing

A total of 1 T1-weighted (T1w) images were found within the input BIDS dataset.The T1-weighted (T1w) image was corrected for intensity non-uniformity (INU) with `N4BiasFieldCorrection` (Tustison et al., 2010), distributed with ANTs 2.3.3 (Avants et al., 2011, RRID: SCR 004757), and used as T1w-reference throughout the workflow. The T1w-reference was then skull-stripped with a *Nipype* implementation of the `antsBrainExtraction.sh` workflow (from ANTs), using OASIS30ANTs as target template. Brain tissue segmentation of cerebrospinal fluid (CSF), white-matter (WM) and gray-matter (GM) was performed on the brain-extracted T1w using `fast` (Smith et al., 2004, 6.0.5.1:57b01774, RRID:SCR 002823). Brain surfaces were recon-structed using `recon-all` (Reuter et al., 2012, FreeSurfer 7.2.0, RRID:SCR 001847), and the brain mask estimated previously was refined with a custom variation of the method to reconcile ANTs-derived and FreeSurfer-derived segmentations of the cortical gray-matter of Mindboggle (Klein et al., 2017, RRID: SCR 002438). Volume-based spatial normalization to one standard space (MNI152NLin2009cAsym) was performed through nonlinear registra-tion with `antsRegistration` (ANTs 2.3.3), using brain-extracted versions of both T1w reference and the T1w template. The following template was selected for spatial normalization: ICBM 152 Nonlinear Asymmetrical template version 2009c (Fonov et al., 2009, RRID: SCR 008796; TemplateFlow ID: MNI152NLin2009cAsym;).

#### Functional data preprocessing

For each of the 3 BOLD runs found per subject (across all tasks and sessions), the following preprocessing was performed. First, a reference volume and its skull-stripped version were generated by aligning and averaging 1 single-band references (SBRefs). Head-motion parameters with respect to the BOLD reference (transformation matrices, and six corresponding rotation and translation parameters) are estimated before any spatiotemporal filtering using `mcflirt` (Jenkinson et al., 2002, FSL 6.0.5.1:57b01774). The estimated *fieldmap* was then aligned with rigid-registration to the target EPI (echo-planar imaging) reference run. The field coefficients were mapped on to the reference EPI using the transform. The BOLD reference was then co-registered to the T1w reference using `bbregister` (FreeSurfer) which implements boundary-based registration (Greve and Fischl, 2009). Co-registration was configured with six degrees of freedom. First, a reference volume and its skull-stripped version were generated using a custom methodology of *fMRIPrep*. Several confounding time-series were calculated based on the *preprocessed BOLD* : framewise displacement (FD), DVARS and three region-wise global signals. FD was computed using two formulations following Power (Power et al., 2014, absolute sum of relative motions,) and Jenkinson (Jenkinson et al., 2002, relative root mean square displacement between affines,). FD and DVARS are calculated for each functional run, both using their implementations in *Nipype* (Power et al., 2014, following the definitions by). The three global signals are extracted within the CSF, the WM, and the whole-brain masks. Additionally, a set of physiological regressors were extracted to allow for component-based noise correction (Behzadi et al., 2007, *CompCor*,). Principal components are estimated after high-pass filtering the *preprocessed BOLD* time-series (using a discrete cosine filter with 128s cut-off) for the two *CompCor* variants: temporal (tCompCor) and anatomical (aCompCor). tCompCor components are then calculated from the top 2% variable voxels within the brain mask. For aCompCor, three probabilistic masks (CSF, WM and combined CSF+WM) are generated in anatomical space. The implementation differs from that of Behzadi et al. in that instead of eroding the masks by 2 pixels on BOLD space, a mask of pixels that likely contain a volume fraction of GM is subtracted from the aCompCor masks. This mask is obtained by dilating a GM mask extracted from the FreeSurfer’s *aseg* segmentation, and it ensures components are not extracted from voxels containing a minimal fraction of GM. Finally, these masks are resampled into BOLD space and binarized by thresholding at 0.99 (as in the original implementation). Components are also calculated separately within the WM and CSF masks. For each CompCor decomposition, the *k* components with the largest singular values are retained, such that the retained components’ time series are sufficient to explain 50 percent of variance across the nuisance mask (CSF, WM, combined, or temporal). The remaining components are dropped from consideration. The head-motion estimates calculated in the correction step were also placed within the corresponding confounds file. The confound time series derived from head motion estimates and global signals were expanded with the inclusion of temporal derivatives and quadratic terms for each (Satterthwaite et al., 2013). Frames that exceeded a threshold of 0.5 mm FD or 1.5 standardized DVARS were annotated as motion outliers. Additional nuisance timeseries are calculated by means of principal components analysis of the signal found within a thin band (*crown*) of voxels around the edge of the brain, as proposed by (Patriat et al., 2017). The BOLD time-series were resampled into standard space, generating a *preprocessed BOLD run in MNI152NLin2009cAsym space*. First, a reference volume and its skull-stripped version were generated using a custom methodology of *fMRIPrep*. All resamplings can be performed with *a single interpolation step* by composing all the pertinent transformations (i.e. head-motion transform matrices, susceptibility distortion correction when available, and co-registrations to anatomical and output spaces). Gridded (volumetric) resamplings were performed using `antsApplyTransforms` (ANTs), configured with Lanczos interpolation to minimize the smoothing effects of other kernels (Lanczos, 1964). Non-gridded (surface) resamplings were performed using `mri vol2surf` (FreeSurfer).

Many internal operations of *fMRIPrep* use *Nilearn* 0.9.1 (Abraham et al., 2014, RRID:SCR 001362), mostly within the functional processing workflow. For more details of the pipeline, see the “FMRIPrep’s documentation” section corresponding to workflows in *fMRIPrep*’s documentation.

### Data analysis: fMRI general linear models

To assess neural correlates of observed and latent variables, we ran whole-brain, GLMs to generate statistical parametric maps at the voxel level using SPM12 (https://www.fil.ion.ucl.ac.uk/spm/software/spm12/ in MATLAB. All GLMs were designed with zero-duration event onset regressors at the times of fixation, subgoal report, response states 1 and 2, responses 1 and 2, response state 2, state 3, and goal presentation. All variables were modeled as parametric modulators of one of these events. Current subgoal and goal amount were locked to response statese 1 and 2, left and right choices were locked to responses 1 and 2, rare (vs. common) transition was locked to response state 2, subgoal collected (vs. not collected) was locked to state 3, SPE was locked to state 3, and indicator variables for encountered outcome (permit, alien, home) were locked to state 3 as well. Current subgoal was not assessed during state 3 to avoid the confound of experienced subgoal and the change in the potential change in current subgoal as a result of the outcome. GLMs only differed by which value-related term was included, which was locked to response states 1 and 2. These GLMs were run separately because of the correlations between value-related terms. For state 1 on rare trials, value-related terms for the inferred option chosen were used. For state 2 on rare trials, value-related terms were computed as a function of action values. For the BMS analysis, GLMs were run without current subgoal as a regressor, since the value-related terms had different amounts of correlation with the variable.

The GLMs also included nuisance regressors, estimated using *fMRIPrep*, to control for signal and signal artifacts unrelated to the task. These regressors included 6 head motion variables (rotation and translation in 3 dimensions), the average signal within the brain mask, framewise displacement (Power et al., 2012), the derivative of RMS variance over voxels (Power et al., 2012), a maximum of 6 anatomical CompCor PCA components, and a maximum of 6 temporal ComCor components (Behzadi et al., 2007).

GlMs were first run for every subject across the 3 functional runs. Contrasts were generated for the variables of interested and were combined to construct second-level T maps using the standard summary statistics approach for random effects analysis. The statistical maps were thresholded at *p <* 0.001 uncorrected, then a minimum cluster size was applied that corresponded to a whole-brain cluster-level, FWE-corrected threshold of *p <* 0.05. The conjunction analysis (Figure 4A) was performed to assess regions commonly activated by subgoals and goals, across the population. In SPM12, we compared the minimum statistic to the conjunction null in order to test for significant conjunction of effects (Nichols et al., 2005). Resulting statistical maps were thresholded at *p <* 0.001 uncorrected and FWE corrected at the cluster level using a *p <* 0.05 threshold.

### Data analysis: fMRI Bayesian Model Selection

BMS was performed using the MACS (Model Assessment, Comparison and Selection) toolbox (Soch and Allefeld, 2018) for SPM in order to determine which value-related decision variables best explained the neuroimaging signal in different regions. Briefly, cross-validated, log model evidence maps were computed for each subject and decision variable GLM at every voxel. Next, cross-validated BMS was used to compare models, voxelwise, at the subject level. At the population level, random-effects BMS analysis was performed to generate exceedance probability maps, defining the likelihood of a given GLM explaining the variance in the neuroimaging signal best, relative to the others. Exceedance probabilities were averaged within ROIs defined by clusters from the GLMs that survived FWE corrections at the cluster level and overlapped with the anatomically-defined region being considered.

### Data analysis: fMRI decoding

We used searchlight MVPA to test for distributed, high-dimensional represen-tations of current subgoal and second action, separately. To extract features from the neuroimaging data, we ran GLMs to estimate betas for each trial, modeled as zero-duration events at the onset of the epochs relevant to the analysis. The same nuisance regressors as those used in the other GLMs described above were included to control for non-task-related variance in the signal. We used the https://nilearn.github.io package in Python to train a linear support vector classifier on these features in spheres of voxels with radius 5 mm. For subgoal decoding, we trained the decoder on activity during the outcome state and tested on data from the first response state. We used the objective current subgoal as the label, given that subjects exhibited reliable understanding of the correct subgoal in the sequence (Figure 1D). For second choice decoding we trained and tested the decoder on activity during the first response state. We limited this decoding to common transition trials and used the observed second action as the label. We decoded second action instead of option because sensory- and motor-related information about the first action was already present in the epoch being assessed, during which the first action was executed. In both cases we used 5-fold stratified cross validation and averaged the generated accuracies over the folds. For subgoal decoding, the subgoal labels were balanced in each fold of the training data (from the outcome state) and all folds were tested on all of the test data (from the first response state). For second action decoding, we found that subjects exhibited option biases: given the same first action, some subjects were more likely to choose one of the two second actions (Figure S5A,B). Because we were analyzing activity during performance of the first action and wanted to avoid the possibility of the decoder leveraging this information, we subsampled the data to equalize the numbers of each type of option that shared the same first action.

Because a second-level *t*-test is inappropriate for information-based measures like decoding accuracy, we ran a very conservative test that leverages the minimum statistic to test for the significant prevalence of information, voxelwise, across the population and controls for multiple comparisons (Allefeld et al., 2016). First, we generated null distributions of whole-brain decoding accuracies for each subject. Labels were randomly permuted and the decoder was fit on the shuffled data using the exact procedure as described above. The null decoders were estimated on 50 shuffled datasets in parallel for each subject using the resources of Caltech’s Resnick High Performance Computing Center in order to run the whole-brain searchlight analyses in a reasonable amount of time. A second-level null distribution was generated by Monte Carlo estimation, randomly sampling a shuffled accuracy for each subject 10, 000 times at each voxel. The minimum accuracy across subjects was found for each voxel. To control for multiple comparisons, instead of minimum, the maximum accuracy was found for each of the 10, 000 null permutations at each voxel. The *p* value of a voxel is the probability that the subjects’ minimum accuracy is greater than the maximum accuracies of all second-level permutations. The threshold for significance was set at *α* = 0.05. For all significant voxels, we then computed the threshold of prevalence (maximum rate of information prevalence across the population) at which the null could be rejected as a function of the *p* value.

---

**Figure S1.**
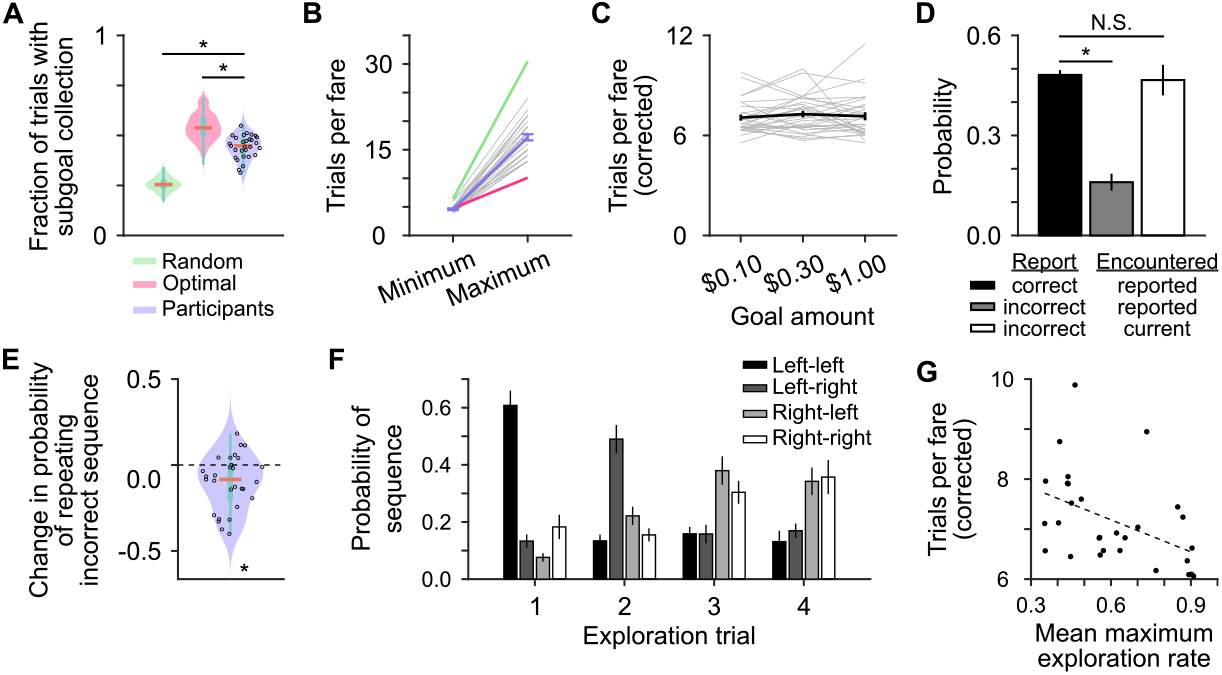
(A) Fraction of trials on which any subgoal was collected (a measure of efficiency correlated with trials per fare) for simulated and actual behavior. Top and bottom edges of blue boxes represent the 75th and 25th percentile, respectively. Orange line is the median of the data. Subject performance was significantly better than the random agent (*W* = 5.0 *×* 10^5^, *p <* 10^*−*19^, Wilcoxon rank sum test) but worse than the optimal agent (*W* = 5.3 *×* 10^5^, *p <* 10^*−*10^, Wilcoxon rank sum test). (B) Minimum and maximum number of trials taken to complete a fare for subjects and simulated agents. Gray lines are individual subjects. (C) There was no effect of fare value on performance (coefficient = 0.040, *t*_85_ = 0.38, *p* = 0.58, linear mixed-effects model). (D) Probability of encountering a subgoal as a function of report, whether the report was correct, and the current subgoal. Subjects were significantly more likely to encounter the correctly-reported subgoal than the incorrectly-reported subgoal (*t*_27_ = 9.67, *p <* 10^*−*9^, paired *t*-test). There was no difference between encountering the correctly-reported subgoal and encountering the current subgoal on incorrect report trials (*t*_27_ = 0.40, *p* = 0.69, paired *t*-test). (E) The probability of repeating a previously-taken, incorrect sequence of actions improved significantly over the session (*t*_28_ = *−*3.44, *p* = 0.0018, one-sample *t*-test). Average probability in last 5 fares - average probability in the first 5 fares, corrected for rare transitions. (F) Average exploration patterns averaged across all subjects. The average reveals a two dominant patterns. First, left-left, followed by left-right, then right-left, and finally right-right. Second, left-left, left-right, right-right, and left-right. (G) Better performance correlates with greater exploration stereotypy (estimate = *−*2.14, *t*_27_ = *−*2.62, *p* = 0.014, linera regression). Average maximum exploration rate is the mean of the highest sequence probabilities of each of the 4 exploration trials. Dashed line shows regression fit.

**Figure S2.**
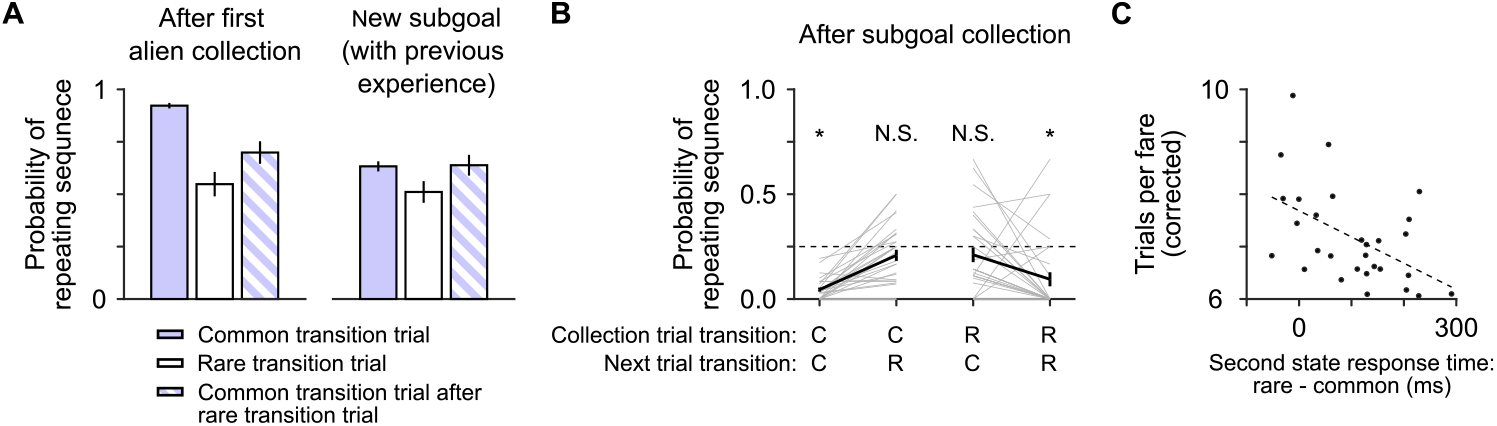
(A) Left: probability of repeating the sequence of actions for acquiring the second alien is affected by forgetting (effect of trial number after first alien, coefficient = *−*0.82, *t*_568_ = *−*5.22, *p <* 10^*−*6^, logistic regression model) and transition type (effect of transition, coefficient = *−*1.47, *t*_568_ = *−*5.19, *p <* 10^*−*6^, logistic regression model). Right: probability of repeating the sequence of actions for acquiring a subgoal that was experienced while irrelevant, during or after a rare transition trial on the first trial in which that subgoal becomes relevant. This probability is significantly lower on the intervening rare transition trial (effect of transition, coefficient = *−*0.51, *t*_588_ = *−*2.16, *p* = 0.03, logistic regression model). Errorbars are Bernoulli SEM. (B) Probability of repeating the same sequence of choices as the previous trial when the current subgoal has changed in between, split by transition type on both trials. “C” = common, “R” = rare. Asterisks indicate *p <* 0.05 from *t*-tests compared to chance (*P* = 0.25). (C) Larger difference in response state 2 response times after rare or common transitions predicts better behavior performance (coefficient = *−*0.0050, *t*_27_ = *−*3.01, *p* = 0.005).

**Figure S3.**
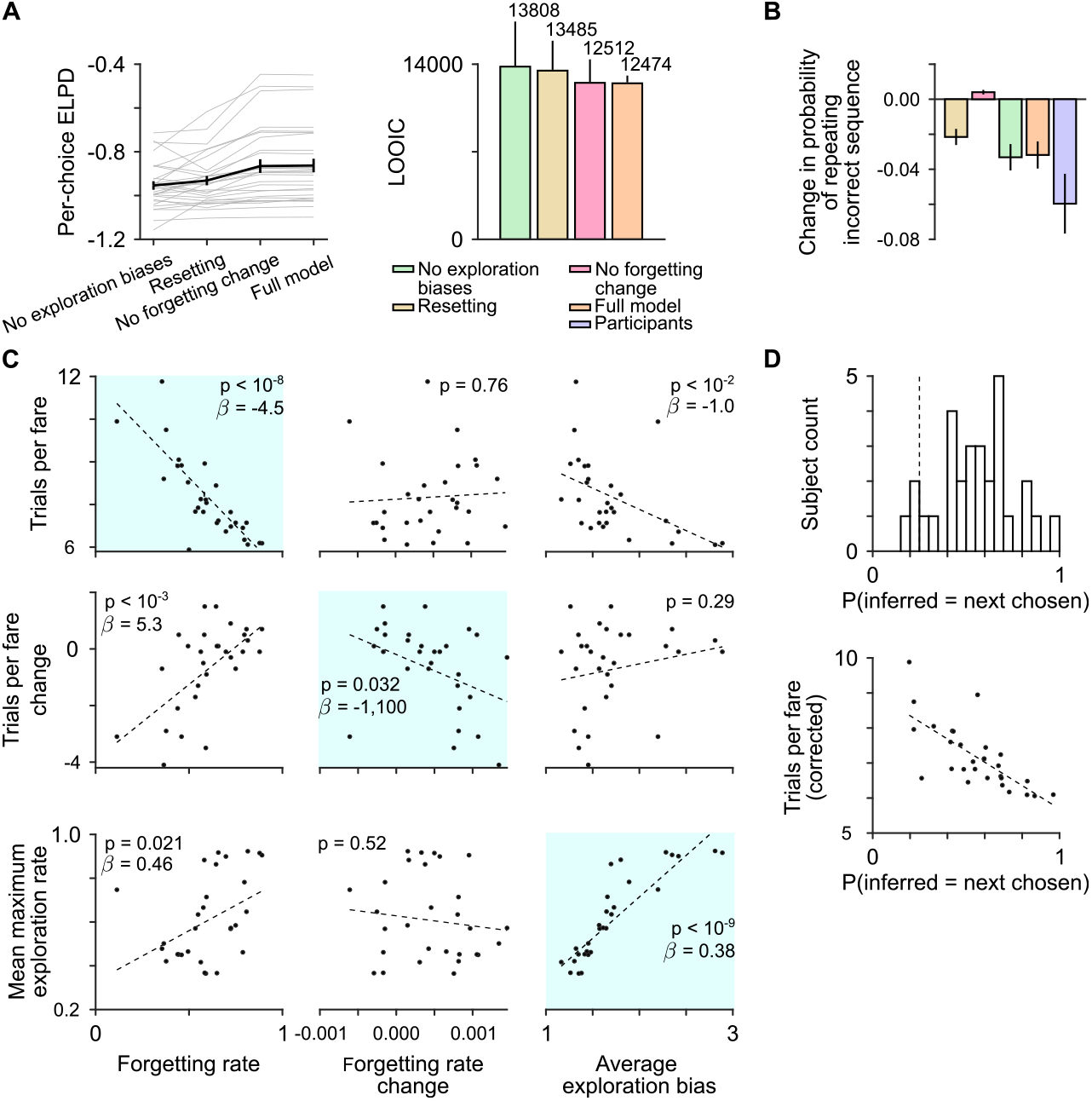
(A) Results from 5-fold cross validation for behavior model comparison. Left: per-choice log predictive densities of held out data. Error bars represent SEM. Right: leave-one-out cross-validation information criterion. (B) Improvement in incorrect behavior over the course of the session, as in Figure S1e, for simulated and participant behavior. Corrected for rare transitions. Colors same as in A. Error bars represent SEM. (C) Comparison of maximum *a posteriori* estimates of parameters from the full MB-HRL model to behavior measures. Light blue squares highlight the features that the parameters were intended to capture. *p*-values from linear models of observed behavior metrics regressed on the behavior model parameters. Regression coefficients (*β*) indicated for significant relationships. (D) Analysis of method for inferring chosen option on rare transition trials. Top: probability that the inferred option matches the option selected on the next common transition trial. Dashed line is chance. Bottom: behavior performance as a function of inferred prediction accuracy.

**Figure S4.**
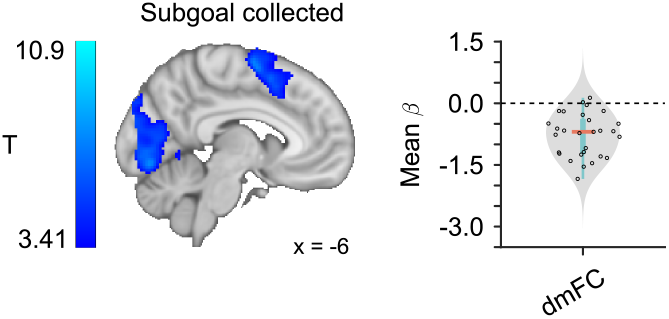
Negative GLM correlations with subgoal collected, tested using a cluster-forming threshold of *p <* 0.001 uncorrected, followed by cluster-level FWE correction at *p <* 0.05. Left: population-level T-statistic maps. Right: individual-level regression coefficients within the significant clusters defined at the population level. Top and bottom edges of blue box represent the 75th and 25th percentile, respectively. Orange line is the median of the data.

**Figure S5.**
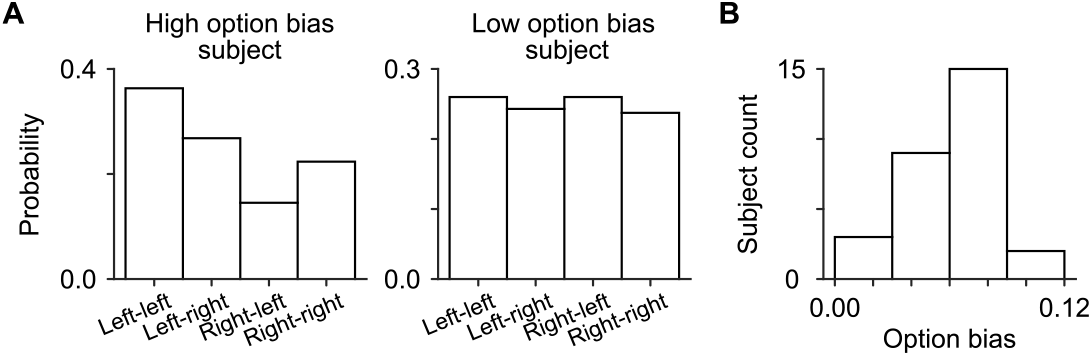
(A) Probability of choosing an option on any trial for example participants with high (left) or low (middle) biases. (B) Distribution of option biases across the population (sum of the absolute value of the differences between pairs of options that begin with the same action).

**Figure S6.**
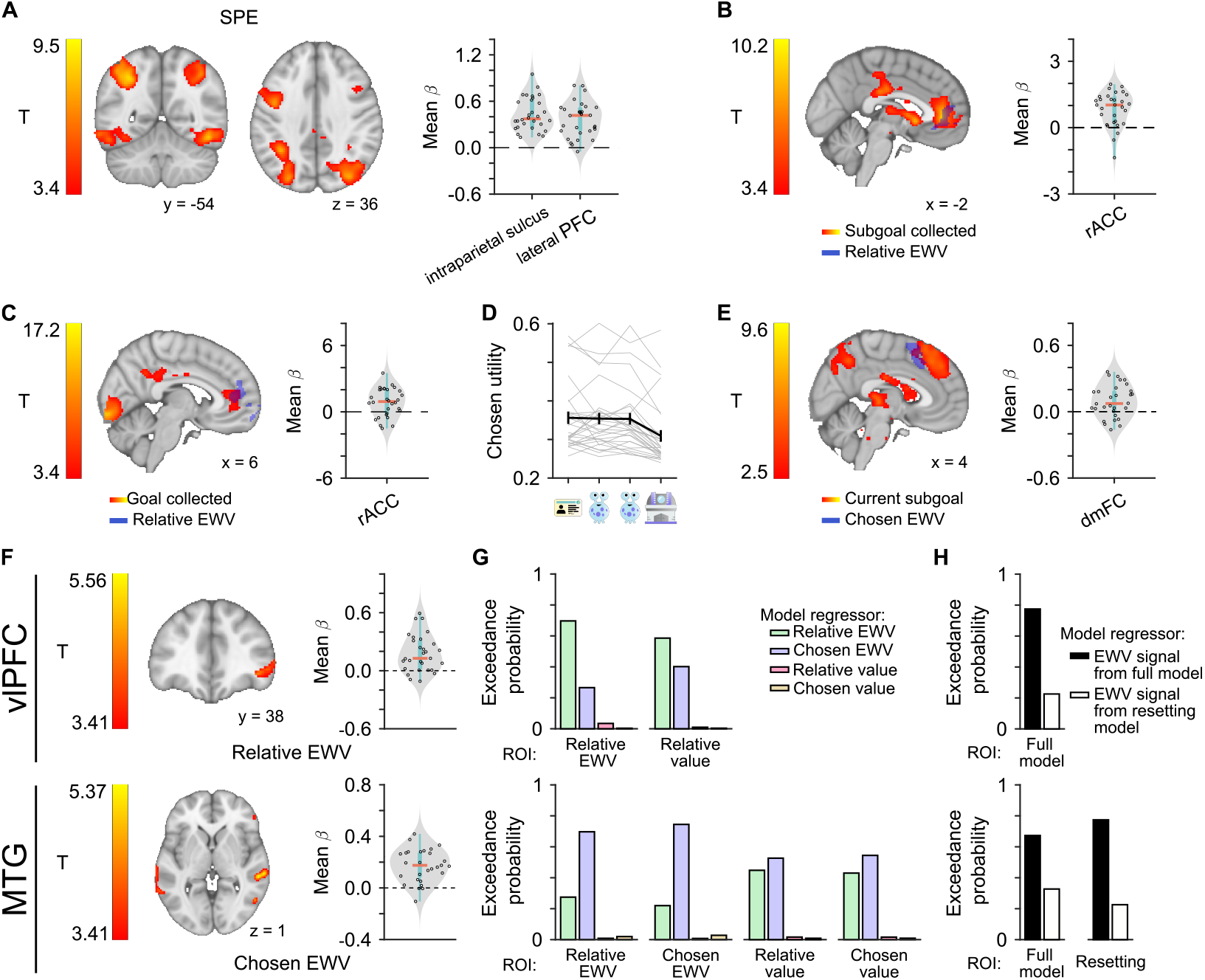
(A) Positive GLM correlations with SPEs tested using a cluster-forming threshold of *p <* 0.001 uncorrected, followed by cluster-level FWE correction at *p <* 0.05. Left: population-level T-statistic maps. Right: individual-level regression coefficients within the significant clusters defined at the population level. Top and bottom edges of blue box represent the 75th and 25th percentile, respectively. Orange line is the median of the data. (B) Left: positive correlations with subgoal collection (FWE-corrected) and negative correlation with relative exploration-weighted value (EWV; binarized cluster from Figure 6a, top panel) from same GLM with relative exploration-weighted value as the value-related variable. Right: subject-level mean regression coefficients for subgoal collection within the ROI defined by the significant relative exploration-weighted value cluster. (C) As in B, except with goal collection. (D) Chosen exploration-weighted value decreases as a function of subgoal in the sequence (coefficient = *−*0.019, *t*_1,14492_ = *−*2.62, *p* = 0.0089, linear mixed-effects model). Gray lines are subject averages. Black line is population average. (E) Left: positive correlations with current subgoal (FWE-corrected) and negative correlation with chosen exploration-weighted value (binarized cluster from Figure 5c) from same GLM with chosen exploration-weighted value as the value-related variable. Right: subject-level mean regression coefficients for current subgoal within the ROI defined by the significant chosen exploration-weighted value cluster. (F) GLM correlations with decision variables tested using a cluster-forming threshold of *p <* 0.001 uncorrected, followed by cluster-level FWE correction at *p <* 0.05. Left: population-level T-statistic maps. Right: individual-level regression coefficients within the significant clusters defined at the population level. Top row: positive correlations between vlPFC and relative exploration-weighted value. Bottom row: negative correlations between dmFC and chosen exploration-weighted value. (G) Results from a BMS comparison of GLMs to test different value-related variables. Each bar plot shows mean exceedance probabilities for the GLMs with different decision variables in an ROI defined by a significant cluster from one of the GLMs. ROIs were only assessed for variables that survived cluster corrections. ROIs were limited to vlPFC (top row), and MTG (bottom row) regions. (H) Results from a BMS comparison of GLMs with (full model) and without (resetting model) latent learning. ROIs were generated in the same fashion as in B.

